# Mechanistic basis for the emergence of EPS1 as a catalyst in plant salicylic acid biosynthesis

**DOI:** 10.1101/2021.08.21.457228

**Authors:** Michael P. Torrens-Spence, Tianjie Li, Ziqi Wang, Christopher M. Glinkerman, Jason O. Matos, Yi Wang, Jing-Ke Weng

**Affiliations:** Whitehead Institute for Biomedical Research, 455 Main Street, Cambridge, Massachusetts 02142, USA; 2Department of Physics, The Chinese University of Hong Kong, Shatin, New Territories, Hong Kong; 3Department of Biology, Massachusetts Institute of Technology, Cambridge, Massachusetts 02139, USA

## Abstract

Unique to plants in the Brassicaceae family, the production of the plant defense hormone salicylic acid (SA) from isochorismate is accelerated by an evolutionarily young isochorismoyl-glutamate pyruvoyl-glutamate lyase, EPS1, which belongs to the BAHD acyltransferase protein family. Here, we report the crystal structures of apo and substrate-analog-bound EPS1 from *Arabidopsis thaliana*. Assisted by microsecond molecular dynamics simulations, we uncover a unique pericyclic rearrangement lyase mechanism facilitated by the active site of EPS1. We reconstitute the isochorismate-derived pathway of SA biosynthesis in *Saccharomyces cerevisiae*, which serves as an *in vivo* platform that helps identify active-site residues critical for EPS1 activity. This study describes the birth of a new catalyst in plant phytohormone biosynthesis by reconfiguring the ancestral active site of a progenitor enzyme to catalyze alternative reaction.

**One sentence summary:** By reconfiguring the active site of a progenitor acyltransferase-fold, EPS1 acquired the unique, evolutionarily new lyase activity that accelerates phytohormone salicylic acid production in Brassicaceae plants.

## MAIN TEXT

Salicylic acid (SA), or 2-hydroxybenzoic acid, is a simple phenolic acid required in higher plants for local and long-distance defense responses following exposure to pathogens (*1*). Additionally, SA also plays roles in photosynthesis, ion uptake and transport, and growth regulation (*2*). Despite the chemical simplicity of SA and its importance in defense signaling, the full scope of SA biosynthetic pathways across the entire plant kingdom have yet to be thoroughly explored.

To date, two major plant SA biosynthetic pathways have been identified downstream of chorismate: the phenylpropanoid-derived pathway (*3*) and the isochorismate-derived pathway (*4*). In the model plant *Arabidopsis thaliana*, a small fraction of SA is produced downstream of phenylalanine ammonia-lyase (PAL) via side-chain shortening of cinnamic acid to benzoic acid, followed by 2-hydroxylation (*5-8*). The majority of SA *de novo* biosynthesized upon pathogen attack is dependent on isochorismate synthase, a plastid-localized enzyme that catalyzes isomerization of chorismate into isochorismate, encoded by the *SALICYLIC ACID INDUCTION DEFICIENT 2* (*SID2*) gene in *A. thaliana (9)* (Fig. 1A). The *A. thaliana* gene *SALICYLIC ACID INDUCTION DEFICIENT 1* (*SID1*) encodes a multidrug and toxin extrusion (MATE) transporter, which is localized in the chloroplast envelope and transports the isochorismate synthesized by SID2 from the plastid to the cytosol (*10-12*). Unlike the bacterial pathway in which SA is produced directly from isochorismate via an isochorismate pyruvate lyase (IPL) (*13*), plants employ a cytosolic GH3 acyl adenylase family enzyme PBS3 that catalyzes regiospecific conjugation of isochorismate with L-glutamate to form isochorismoyl-glutamate A (IGA) (*12, 14*) (Fig. 1A). IGA is unstable and decays spontaneously into the phytohormone SA and the byproduct *N*-pyruvoyl-L-glutamate (NPG) (*12, 14*). Unique to the Brassicaceae plants including *A. thaliana*, this spontaneous decay reaction is accelerated by a lineage-specific isochorismoyl-glutamate pyruvoyl-glutamate lyase (IPGL) encoded by the *ENHANCED PSEUDOMONAS SUSCEPTIBILTY 1* (*EPS1*) gene (*14*) (Fig. 1A). The *A. thaliana eps1* mutant is hypersusceptible to the bacterial pathogen *Pseudomonas syringae*, and is defective in *de novo* SA production upon *P. syringae* infection (*14, 15*), suggesting the critical function of EPS1 in defense signaling in Arabidopsis. How Brassicaceae plants gained and integrated a newly evolved IPGL in their SA biosynthetic pathway remains unknown.

**Figure 1.**
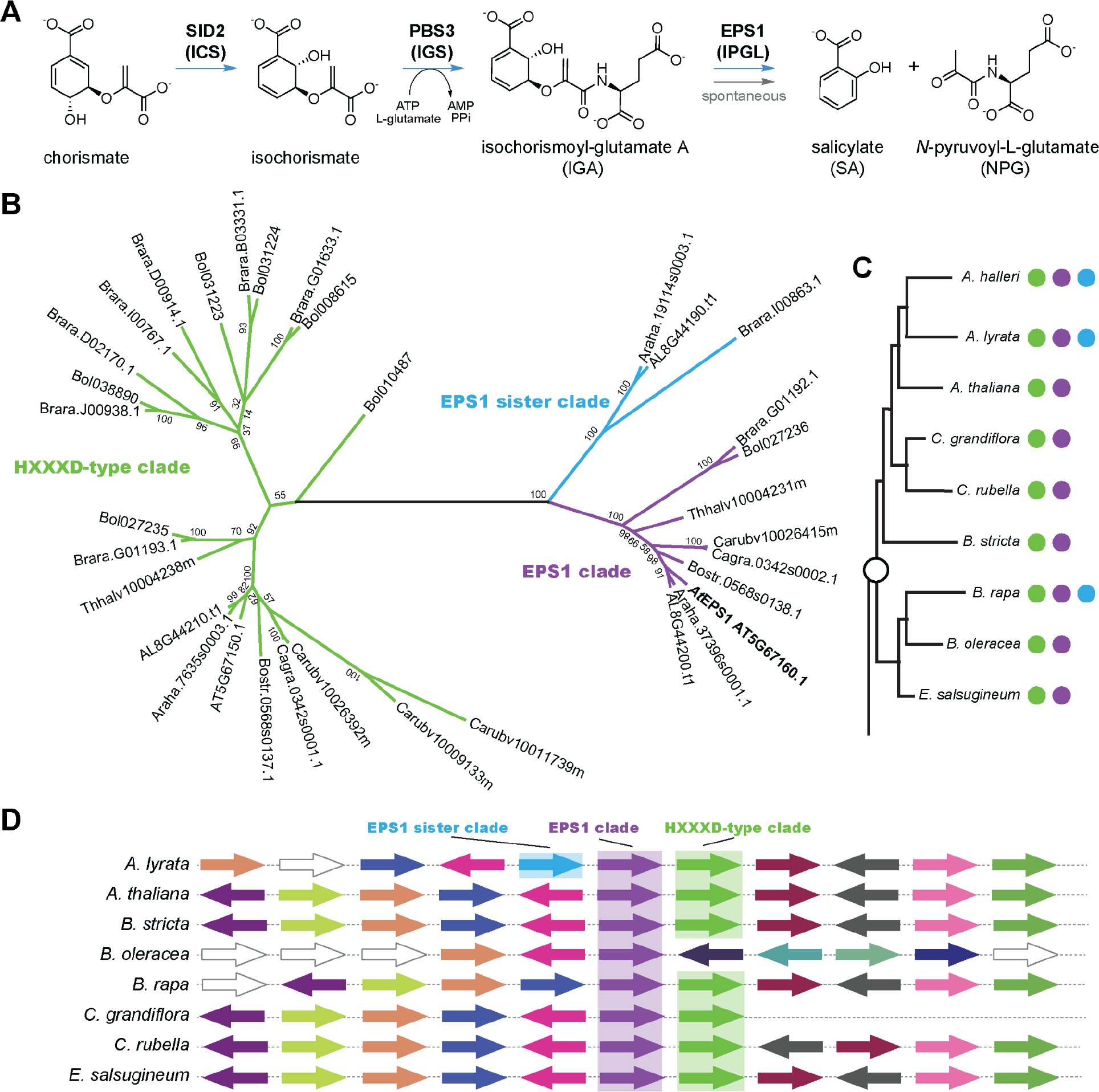
The metabolic function and evolutionary origin of EPS1. (**A**) The SA biosynthetic pathway in Brassicaceae plants. (**B**) Simplified maximum likelihood tree of the EPS1 clade (purple), the EPS1 sister clade (blue) and its most closely related HXXXD-type acyl-transferase family protein ancestral clade (green). Note, the blue EPS1 sister clade is lost in the majority of Brassicaceae plants. The purple EPS1 clade enzymes and not the neighboring clades display strict conservation for the serine substitution for the characteristic histidine in the (HXXXD) motif. (**C**) Simplified taxonomy of Brassicaceae plants displaying the presence or absence of the green ancestor clade, the blue EPS1 sister clade or the purple true EPS1 clade enzymes. (**D**) In *A. thaliana*, the EPS1 gene (purple arrow and shading) is adjacent to its closest homolog AT5G67150.1 (green arrow and shading). In the related species Arabidopsis lyrata, EPS1 is sandwiched between the AT5G67150.1 homolog AL8G44210.t1 (green clade) and the even more closely related *A. lyrata* AL8G44190.t1 (blue arrow and shading). Analysis suggests that in *A. thaliana*, AT5G67150.1 duplicated to produce a AL8G44190.t1 homolog which duplicated again to make EPS1. The AL8G44190.t1 homolog was then lost in *A. thaliana* and the majority of other profiled Brassicaceae species.

EPS1 is a member of the plant BAHD acyltransferase family, which is named after the first four enzymes characterized within the family (*16*). The BAHD family has undergone extensive radiation during land plant evolution, giving rise to functionally diverse acyltransferases widely distributed in many plant natural product biosynthetic pathways (*17*). Canonical BAHD enzymes catalyze the transfer of the acyl group from an acyl-CoA thioester substrate to an alcohol or amine-containing acceptor molecule to form the corresponding ester or amide product. Most BAHD enzymes contain a conserved catalytic histidine, which serves as the general base that deprotonates the -OH or -NH_2_ group of the acyl acceptor substrate to initiate the catalytic cycle (*18*). It was noted that *A. thaliana* EPS1 (*At*EPS1) and its orthologs found in several other Brassicaceae species were the only BAHD-family proteins identified to date that contain a serine substitution at this conserved histidine residue (*14*), implicating a reconfigured catalytic machinery adapted for the neofunctionalized IPGL activity in EPS1.

To retrace the evolutionary history of EPS1, we first performed a focused phylogenetic analysis of 62 annotated BAHDs encoded by the *A. thaliana* genome. *At*EPS1 was found to belong to the Ib clade, where select clade I members were previously reported to be involved in flavonoid biosynthesis (Fig. S1) (*16, 19*). Expanded phylogenetic analysis of additional EPS1-like homologs from multiple reference plant genomes further revealed that EPS1 falls into a subgroup of BAHDs within the Ib clade that are restricted to Brassicaceae species (Fig. 1B-C). Within this subgroup, the HXXXD-type clade (green, represented by AT5g67150 from *A. thaliana*) is present in all Brassicaceae species examined, and sequences within this clade contain the conserved catalytic histidine indicative of canonical acyltransferase functions. This HXXXD-type clade most likely was the immediate ancestral clade from which the EPS1 clade (purple) and the EPS1 sister clade (blue) were derived (Fig. 1B). Synteny analysis of *EPS1*-containing genomic loci across several Brassicaceae plant genomes provided further details regarding the evolutionary birth of *EPS1* (Fig. 1D). In the *A. thaliana* genome, *AtEPS1* is adjacent to its most closely related homolog *AT5G67150* of the HXXXD-type clade on chromosome 5, which is also the case in the syntenic regions for 7 of the 8 Brassicaceae species examined (Fig. 1D). Moreover, the syntenic region in *A. lyrata* genome harbors the third homologous gene *AL8G44190* of the EPS1 sister clade on the other side of *AlEPS1* (Fig. 1D), capturing the historical founding tandem gene duplication event that ultimately gave rise to *EPS1*, as well as the EPS1 sister clade gene which was subsequently lost in *A. thaliana* and the majority of other profiled Brassicaceae species (Fig. 1C).

To understand the molecular basis underlining how the unprecedented IPGL activity could arise on the basis of a BAHD-fold progenitor, we solved the crystal structures of apo *At*EPS1 and its complex with an unreactive substrate analog (2-(3-carboxyphenoxy)acetyl)-L-glutamic acid (CAG) at 1.9 Å and 1.7 Å resolution, respectively (Fig. 2A, B, Fig. S2-8). Similar to several previously reported BAHD structures, *At*EPS1 displays pseudo-symmetric N-terminal (residues 1-181) and C-terminal (residues 227-434) domains, connected by a large crossover loop (residues 182-226) which runs around ¾ of the circumference of the protein (*18, 20–26*) (Fig. 2A, C). Like other BAHD acyltransferases, the *At*EPS1 active site where CAG binds is located at the interface between the two pseudo-symmetric domains (Fig. 2A). Interestingly, when superpositioning CAG-bound *At*EPS1 with the product-bound structure of *A. thaliana* hydroxycinnamoyl-CoA:shikimate hydroxycinnamoyltransferase (*At*HCT) (PDB ID: 5KJU) (*18*), a canonical BAHD acyltransferase, it is apparent that the CAG-binding site in *At*EPS1 corresponds to only the acyl acceptor portion of the *At*HCT active site, whereas several bulky residues in *At*EPS1, notably Leu42, Phe156, Tyr158, Ile168, and Trp169, fill the space otherwise would host acyl donor binding in a canonical BAHD acyltransferase (Fig. 2C). In addition to the substitution of the conserved catalytic histidine to serine in *At*EPS1, Trp371 (*At*HCT numbering), a residue that serves as an oxyanion hole to stabilize the tetrahedral intermediate of the acylation reaction, conserved in a large fraction of BAHD acyltransferases (*18*), is substituted to serine (Ser367) in *At*EPS1 (Fig. 2C).

**Figure 2.**
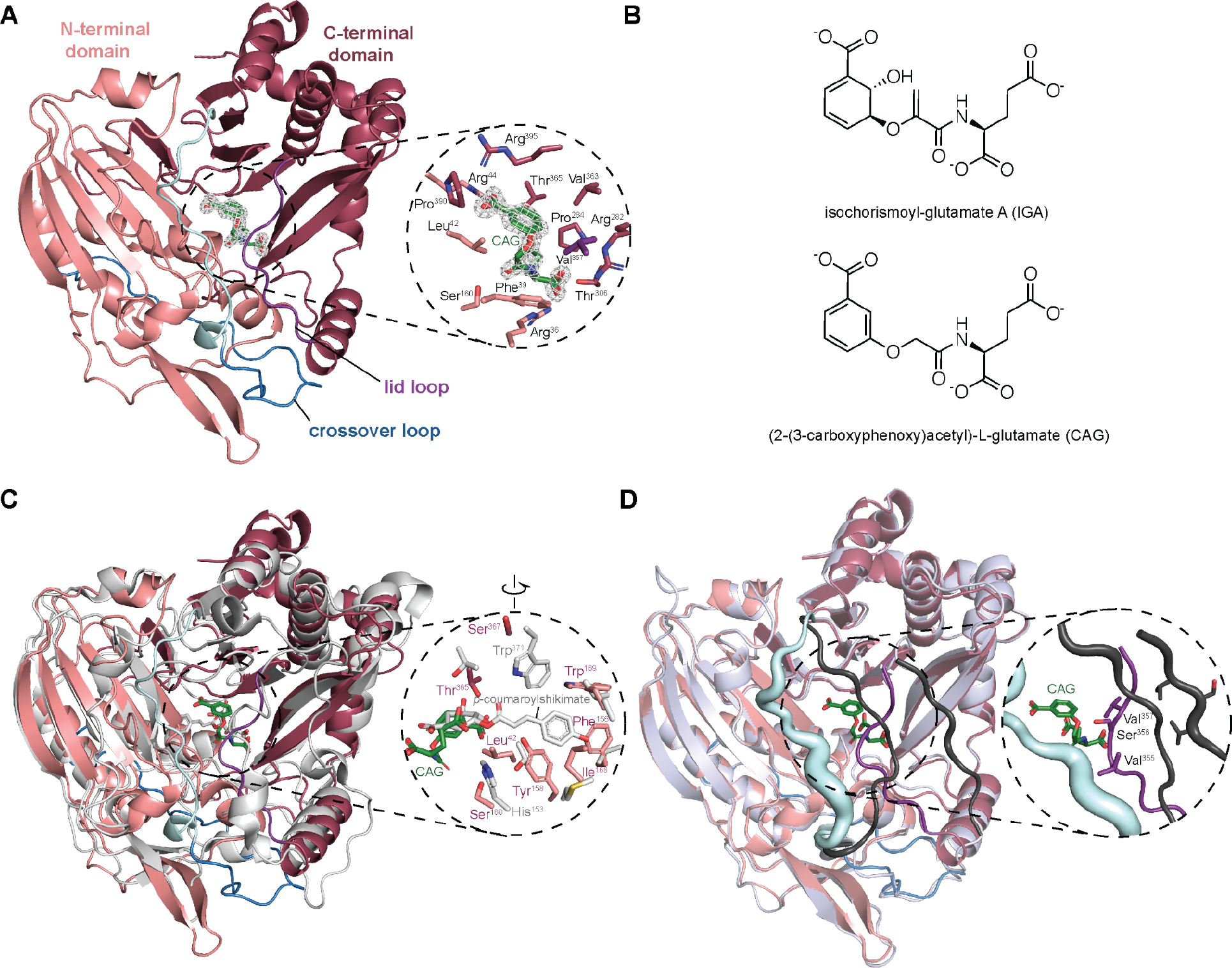
Structural features of *At*EPS1. (**A**) The CAG-bound *At*EPS1 structure rendered as a cartoon. The N-terminal domain is displayed in salmon, the crossover-loop structure is displayed in slate blue, the dynamic portion of the crossover-loop region is colored in cyan, the C-terminal domain is displayed in maroon, and the lid-loop is rendered in purple. |2Fo – Fc| electron density map for the green CAG ligand is contoured at 1.5 σ. (**B**) Structure of the EPS1 ligand (CAG) and the native substrate (IGA). (**C**) The CAG-bound *At*EPS1 and the grey superimposed *p*-coumaroylshikimate-bound *At*HCT (PDB: 5KJU). *At*EPS1 residues corresponding to the *At*HCT catalytic triad and the bulky residues filling the acetyl donor site are individually labeled in salmon. The two catalytic residues of *At*HCT are labeled in grey. (**D**) Superimposition of the *At*EPS1 CAG-bound and the apo structures. The apo structure is displayed in bluewhite with the dynamic loops rendered in black. The dynamic loops from both structures are displayed in cartoon putty where the radius of the loops represents relative b-factor. Val355, Ser356, and Val357, annotated in the CAG-bound model, are the only dynamic-loop residues in either structures, within 6 Å distance from CAG.

The active site of EPS1 exhibits features that support specific substrate binding (Fig. 2A). The aryl carboxylate of CAG is coordinated through the backbone carbonyl of Pro390, in addition to the positively charged guanidino groups of Arg44 and Arg395. The aromatic ring of CAG is stabilized by the side chains of Leu42, Val357, Val363, and Leu392, while Phe39 likely functions as part of this hydrophobic pocket to hold the acetyl portion of CAG or the corresponding acrylate group of the true IGA substrate. The ɑ-carboxyl group of the glutamate portion of CAG is highly coordinated by the guanidino groups of Arg36 and Arg282 as well as the hydroxyl group of Thr306, while the γ-carboxyl group of the glutamate portion forms a H-bond with the hydroxyl group of Thr365.

Amongst previously reported plant BAHD structures, the size and arrangement of the crossover loop on the acyl-donor side of the enzyme vary little between structures, while portions of the crossover loop on the acyl-acceptor side of the enzyme vary substantially corresponding to individual enzymes’ acyl-acceptor substrates. Despite its overall high sequence and structural similarity with *At*HCT, *At*EPS1 adopts a crossover-loop conformation more similar to that observed in the vinorine synthase and anthocyanin malonyltransferase structures (*26, 27*), indicating a possible preferred configuration for larger substrates, such as IGA, vinorine, or anthocyanin, compared to the smaller HCT acyl acceptor substrate shikimate (Fig. S9 and Sup. Table S2 and S2). Comparison of the apo- and CAG-bound *At*EPS1 structures reveals conformational changes of a dynamic portion of the crossover loop (Pro212-Asn226), a phenomenon which was not observed in other BAHD structures (Fig. 2D). In the CAG-bound *At*EPS1 structure, crossover-loop residues Pro212-Asn226 are rotated away from the central ligand-binding cavity, and an adjacent dynamic loop (Ile351-Lys360), recently described as the lid loop (*28*), is positioned directly above the substrate analog. Conversely, in the apo *At*EPS1 structure, the lid loop (Ile351-Lys360) and the crossover-loop residues Pro212-Asn226 are displaced to open up a larger entrance to the enzyme active site (Fig. 2D). We employed molecular dynamics (MD) simulations (Sup. Table S4) to investigate the apparent coordinated movement of these active-site loops with ligand binding. Clustering analysis of altogether 16-μs trajectories revealed markedly reduced flexibility of both loops upon EPS1’s transition from its apo to the holo state (Fig. S10A). Between the two dynamic loops, the crossover loop appears to be more tightly locked upon such transitions, while the lid loop retains greater flexibility in the holo state (Fig. S10B). These changes in loop dynamics may contribute to selective substrate binding of EPS1.

Guided by the CAG positioning in the *At*EPS1 active site, we used MD simulations to determine the binding pose of the native substrate IGA. Simulated CAG and IGA binding to *At*EPS1 both conform to a nearly identical binding position compared to the substrate analog in the CAG-bound *At*EPS1 crystal structure, where their differences derive from the non-planar isochorismate ring and the presence of an additional alkene double bond in the native substrate IGA (Fig. 3A, Fig. S11-12). Here, the refined binding pose of the simulated IGA is well retained between altogether 8-μs plain MD as well as simulated annealing calculations, and therefore represents the probable binding configuration with a high degree of confidence. Interestingly, comparison of simulated IGA in *At*EPS1 holo complex and in water reveals that IGA adopts a similar conformation in both, with only the D2 distance between the ether oxygen and the amide hydrogen showing a recognizable decrease in interatomic distance in the enzyme context (Fig. 3B). While the decrease in D2 distance of IGA when positioned in the *At*EPS1 active site could support a E1 elimination enzyme mechanism like that previously proposed for the nonenzymatic decay of IGA (*12*), the long D3 distance (> 4 Å) would likely preclude the deprotonation of the ring hydrogen by the amide to complete the proposed C-O bond cleavage. Meanwhile, simulations of chorismoyl-glutamate A (CGA), an isomer of IGA, illustrate a different orientation of the ring from the native substrate, confirming that CGA serves as a poor substrate for EPS1 (Fig. S13)(*14*). To further explore the catalytic mechanism of IPGL, we next simulated *At*EPS1 bound to its products SA and NPG. These simulations indicate that EPS1-mediated IGA decomposition to form SA and NPG results in a 1.4 Å translation and 57° rotation of the SA ring in addition to a 2.4 Å shift of the oxygen atom in the pyruvoyl moiety of NPG compared to the modeled IGA substrate (Fig. 3C). These apparent conformational changes of the products compared to the substrate suggest that the enzyme imposes an energetically unfavorable conformation of the native substrate IGA. Aromatization alters the pseudodiaxial conformation, elongating the C^3^-O distance, placing strain on the IGA substrate to break the C-O bond, and steering the separation of the products through a pericyclic mechanism (Fig. 3C, D) (*29*).

**Figure 3.**
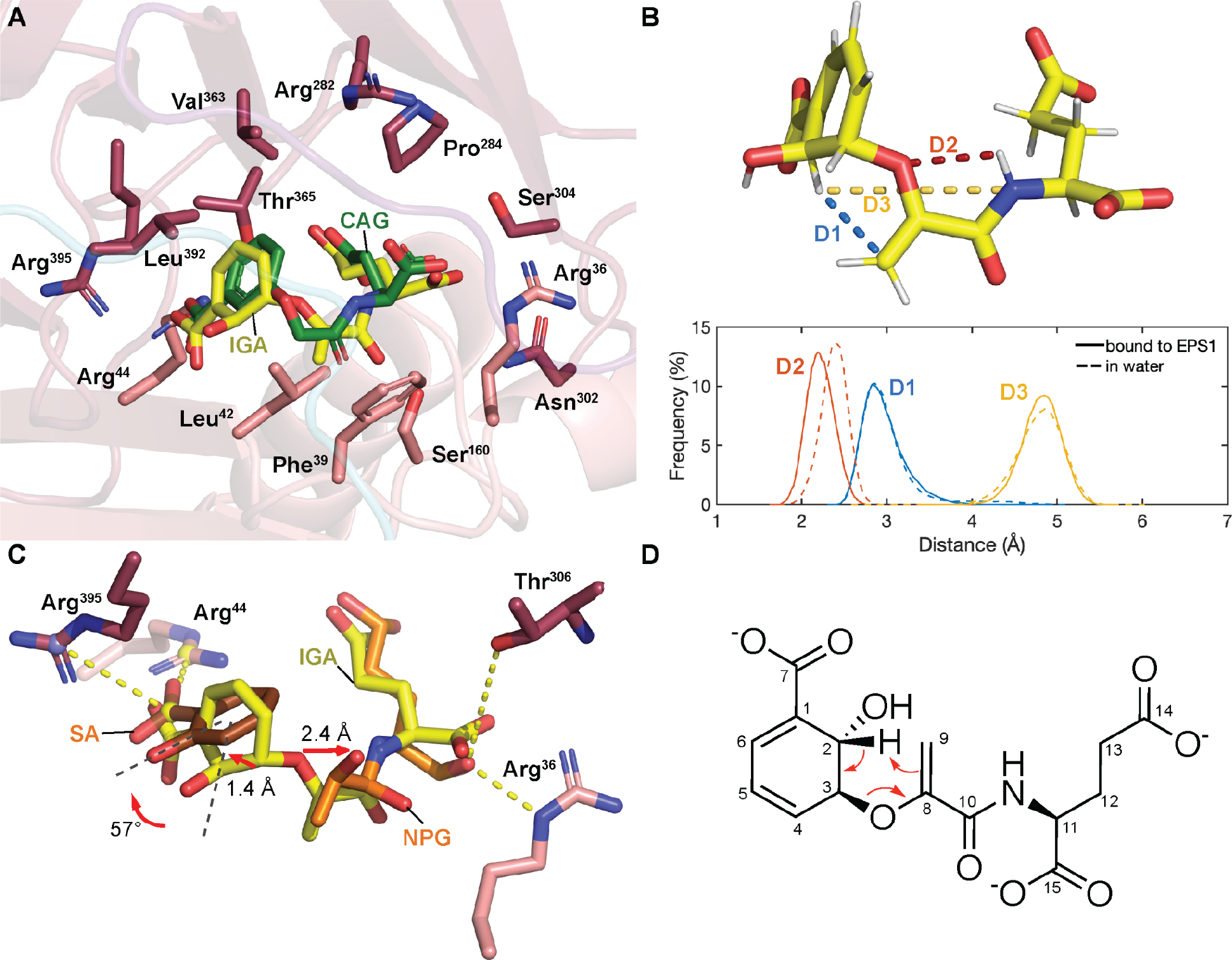
The catalytic mechanism of EPS1. (**A**) IGA rendered in yellow bound to the EPS1 active pocket revealed by altogether 8-μs MD simulations. Crystal structure of its stable analog (2-(3-carboxyphenoxy)acetyl)-L-glutamate (CAG) is shown in green. (**B**) Structural difference of IGA in water (dashed line) and when bound to EPS1 (solid line) measured by three representative distances: D1:H^2^-C^9^ (blue), D2: ether O-amide H (orange), and D3: H^2^-N (green). (**C**) IPL process of IGA in EPS1. Rearrangement results in intramolecular strain release and separation of products through a 57° flipping of SA adjusted by two residues (Arg^395^ and Arg^44^) and anchoring of NPG with Thr^306^, leading to an irreversible IPL process. (**D**) Proposed pericyclic reaction where the hydrogen atom at C^2^ is transferred to C^9^ of the side chain simultaneously with C-O cleavage.

Previously, we characterized the biochemical function of EPS1 *in vitro* using a pre-assay that enzymatically synthesizes the EPS1 substrate IGA from chorismate using recombinant SID2 and PBS3 (*14*). However, the unstable nature of several intermediates, including chorismate, isochorismate, and IGA, precludes the utility of this setup to quantitatively assess the relative activities of site-directed EPS1 mutants. Thus, we sought to reconstitute the isochorismate-derived SA biosynthetic pathway in transgenic *Saccharomyces cerevisiae*, which would allow us to measure relative activities of EPS1 mutants *in vivo*. When experimenting with various combinations of *A. thaliana* genes that enable SA production in *S. cerevisiae*, we discovered that transgenic co-expression of *SID2* and *PBS3* alone is sufficient to elicit SA production in yeast, while the addition of *AtEPS1* further led to seven- and six-fold increase in SA and NPG production, respectively, with a concomitant depletion of IGA levels by more than ten fold (Fig. 4A-C). Although the isochorismate-derived SA biosynthetic pathway is naturally compartmentalized in plastid and cytoplasm in plants, it can be reconstituted in the yeast cytosol without the need for the MATE transporter SID1.

**Figure 4.**
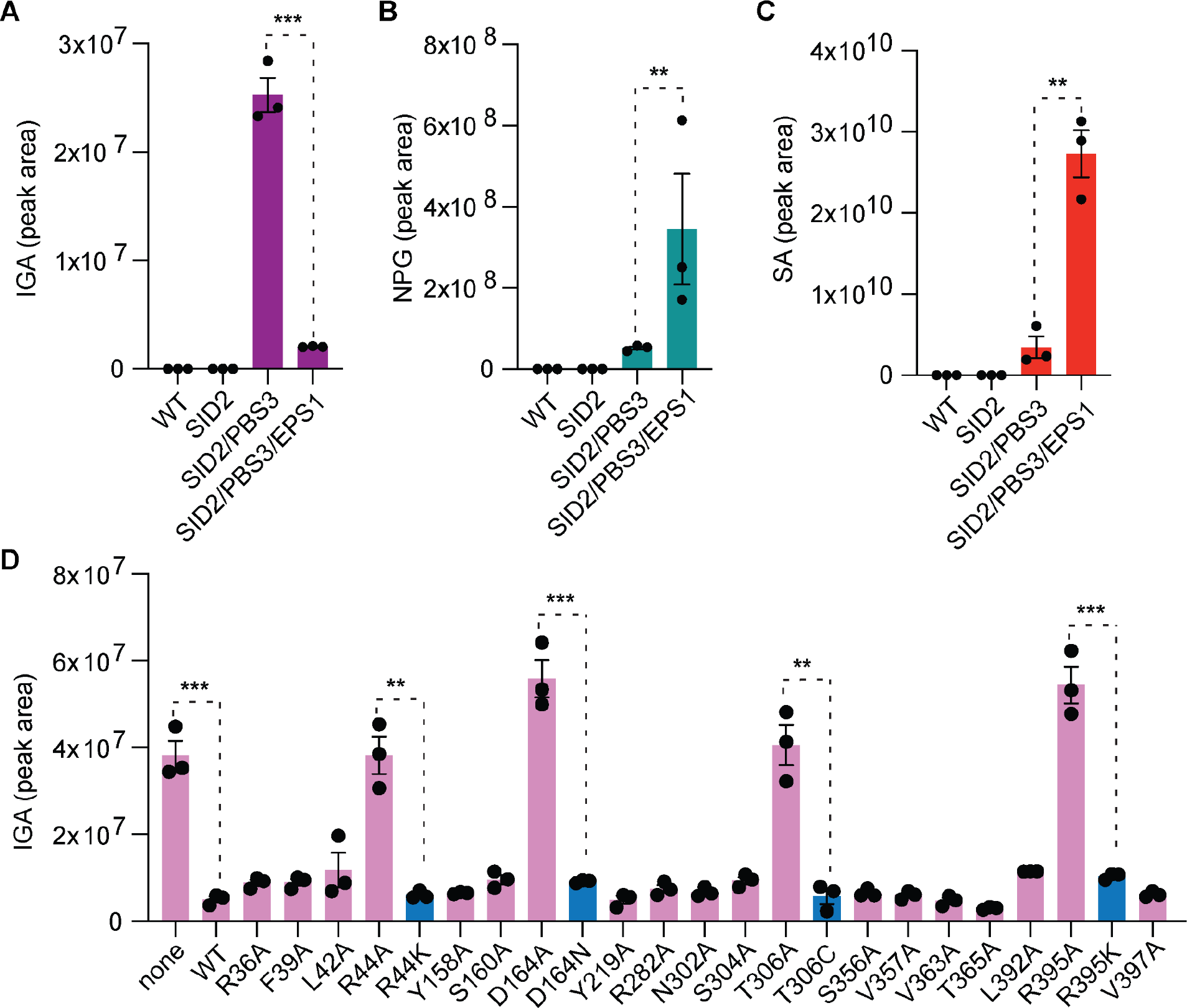
Relative SA pathway metabolite accumulation in various transgenic yeast lines. Error bars indicate standard error of the mean (SEM) based on biological triplicates. Statistical analysis was conducted by two-tailed unpaired t-test. *P < 0.05, **P < 0.01, and ***P < 0.001. (**A**) IGA. (**B**) SA. (**C**) NPG. (**D**) Relative depletion of IGA in *At*EPS1 alanine mutants alongside rescuing mimetic mutations (blue bars) in the heterologous yeast system which additionally express SID2 and PBS3.

Using this newly developed yeast system, we examined the role of various active-site residues of *At*EPS1 by site-directed mutagenesis, followed by *in vivo* IPGL activity measurement. A panel of *At*EPS1 mutants carrying single alanine substitutions of 19 active-site residues were generated and introduced to an *S. cerevisiae* background strain that co-expresses *SID2* and *PBS3*. The depletion of IGA relative to that of the control experiment using wild-type *At*EPS1 was used to quantify the relative IPGL activities of *At*EPS1 mutants (Fig. 4D). Alanine substitution, but not lysine substitution, of Arg44 or Arg395 significantly hampers the IPGL activity, which corroborates the purported roles of Arg44 or Arg395 in coordinating the aryl carboxylate group of IGA through salt bridges and facilitating the leaving of SA product upon pericyclic reaction. Moreover, whereas the Thr306Ala mutation abolishes the IPGL activity in *At*EPS1, the Thr306Cys mutant retains wild-type activity, which verifies the critical role of Thr306 in coordinating the glutamate ɑ-carbonyl group of IGA through hydrogen bonding. The Asp164Ala mutant shows a significantly reduced activity which could be restored by an asparagine substitution. The R group of Asp164 points away from the active site and interacts with Arg288, therefore, Asp164 may play an important role in maintaining the structural integrity of *At*EPS1. Sequence conservation analysis of the active-site residues among EPS1-clade enzymes and EPS1-sister-clade enzymes demonstrates that while Arg44 and Asp164 are required for IPGL activity, these residues are also conserved in the EPS1 sister clade and therefore may not indicate adaption specific for EPS1 neofunctionalization (Fig. S14). Thr306 and Arg395, however, are conserved among the EPS1-clade sequences, but are substituted to other amino acids in the EPS1 sister clade, suggesting these two residues are selected to support IPGL activity among EPS1 orthologs.

The ability to mount a vigorous defense against invading pathogens at local tissue while sending warning signals to the unaffected distal tissue is critical for plants to survive the challenging biotic environments they live in. Although the defense hormone SA could be produced through spontaneous decay of IGA in plants that contain PBS3, natural selection has propelled the common ancestors of the Brassicaceae plants to recruit a gene-duplication-derived BAHD progenitor enzyme, refurbish its active site, and ultimately gain specific IPGL activity that accelerates this last step of SA biosynthesis. In this rare case of neofunctionalized catalytic machinery, EPS1 developed a highly coordinated substrate-binding site that forces an energetically unfavorable conformation for its substrate IGA, thereby facilitating the breakdown of IGA to yield SA and NPG through a pericyclic reaction (Fig. 3D). While pericyclic reactions are regularly used in organic chemistry, enzymes known to catalyze pericyclic reactions have only been sparsely characterized in cellular metabolism, including IPL and chorismate mutase (*30, 31*). Both enzymes employ a pericyclic dissociative mechanism to cleave a C−O bond, which points to a broader mechanism of pericyclic elimination from shikimate-derived metabolites. A systematic understanding of the structural, catalytic, and evolutionary mechanisms underlying these enzymes will ultimately enable us to build future designer catalysts that fulfil diverse chemical engineering and bioengineering needs.

## Acknowledgements

We thank the Metabolite Profiling Core Facilities and the Bioinformatics and Research Computing of the Whitehead Institute for help with the metabolomics experiments and data analysis. This work was supported by the Pew Scholar Program in the Biomedical Sciences (J.K.W.), the Searle Scholars Program (J.K.W.), Keck Foundation (J.K.W.), the National Science Foundation (CHE-1709616, J.K.W.), and project 14306819 from the Hong Kong research grants council (Y.W.). We thank Pingtao Ding for constructive comments on this manuscript.

## Author Contributions

M.P.T.S. and J.K.W. designed the research. M.P.T.S., C.M.G., and J.O.M. performed the cloning and metabolomics sample preparation. C.M.G. performed chemical synthesis. M.P.T.S. and J.K.W. generated and processed the crystallographic data. T.L., Z.W. and Y.W. conducted and analyzed the molecular dynamics simulations. M.P.T.S., C.M.G., T.L., Y.W. and J.K.W. interpreted the results and wrote the paper.

## Supplementary Information

### METHODS

#### Sequence alignment, phylogenetic analysis and homology modeling

Sequence alignments were built using ClustalW2 with default settings (*1*). Maximum likelihood tree shown in Supplementary Fig. 1B and S1 were determined using MEGAX (*2*). The bootstrap consensus trees were inferred from 500 replicates to represent the local BAHD phylogeny of AtEPS1 which encompassed homologous sequences from the phytozome SMB clade with an e-value threshold of 1e-40. The local EPS1 tree was then generated as described above from only the EPS1 clade sequences and a probable sister and ancestor clade sequences. Conservation of the active-site residues between the EPS1 and probable sister clade was displayed using WebLogo (*3*).

#### Molecular cloning, heterologous expression, and recombinant protein production

Total RNA from *A. thaliana* was extracted from six week old plants grown in long-day greenhouse conditions using the RNeasy Mini Kit (Qiagen). First-strand cDNAs were synthesized using the Invitrogen SuperScript III kit (Invitrogen) with the oligo(dT)20 primer. The coding sequences (CDS) of candidate genes were amplified from cDNAs by PCR using gene-specific primers. Point mutations were produced with gibson assembly mutagenesis using the initial cloning primers and the mutation primers. The open reading frame (ORF) of target SA biosynthetic genes were cloned into p423TEF, p425TEF, and p426TEF 2μ plasmids for constitutive expression in *S. cerevisiae* or into pHis8-4, a bacterial expression vector containing an N-terminal 8xHis tag followed by a tobacco etch virus (TEV) cleavage site for recombinant protein production in E. coli. Note, recombinant protein expression of *At*EPS1 was performed as previously described (*4*). 15 mL cultures of transgenic S. cerevisiae BY4743 strains were grown in 50 mL mini bioreactor tubes for 24 hours with shaking at 30 °C. The cultured cells were subsequently pelleted, washed, disrupted, and clarified for LC-HRAM-MS metabolic profiling as previously described (*5*). The raw data were processed using MZmine 2 (*6*) and further analyzed using metaboanalyst (*7*). Statistical analysis was conducted using Prism 8 (GraphPad Prism version 8.0.0 for Mac, GraphPad Software, San Diego, California USA, www.graphpad.com).

#### Chemical synthesis General Methods

All reactions were performed under nitrogen unless otherwise noted. All reagents and solvents were used as supplied without further purification unless otherwise noted. Column chromatography was conducted using Silicycle SiliaFlash P60 SiO_2_ (40–63 μm). Analytical TLC was conducted using Millipore SiO_2_ 60 F_254_ TLC (0.250 mm) plates. HPLC was conducted using a pair of Shimadzu LC-20AP pumps, a Shimadzu CBM-20A communications module, a Shimadzu SPD-20A UV-Vis detector, and a Phenomenex Kinetex 5u C18 100 Å Axia column (150 x 21.2 mm). Melting points were obtained using a Mel-Temp II apparatus and are uncorrected. ^1^H and ^13^C NMR spectra were obtained using a Bruker Avance Neo 400 MHz spectrometer equipped with a 5 mm Bruker SmartProbe. IR spectra were obtained using a Bruker Alpha 2 with a Platinum ATR accessory. Mass spectrometric analysis was performed on a JEOL AccuTOF-DART. Syntheses were inspired by previously described approaches towards polymer-supported IBX (*8*).

#### Protein Crystallization and Structural Determination

Crystals for *At*EPS1 and (2-(3-carboxyphenoxy)acetyl)-L-glutamic acid bound *At*EPS1 were grown at 4 °C by hanging-drop vapor diffusion method with the drop containing 0.9 µL of protein sample and 0.9 µL of reservoir solution at a reservoir solution volume of 500 µL. The crystallization buffer for the both *At*EPS1 structures were composed of 0.2 M Potassium Sodium Tartrate 20 % (w/v) PEG 3350. The ligand bound *At*EPS1 drop also contained 500µM (2-(3-carboxyphenoxy)acetyl)-*L*-glutamic acid. Crystals were cryogenized with an additional 15% weight/volume ethylene glycol. The structures were determined first by molecular replacement using the native HCT structure from *Coffea canephora* (PDB:4G0B) (*9*) as the search model in Molrep (*10*). The resulting model was iteratively refined using Refmac 5.2 (*11*) and then manually refined in Coot 0.7.1 (*12*).

#### MD Simulation

Molecular dynamics simulations of apo- and CAG-bound *At*EPS1 were constructed using the corresponding crystal structures. For holo-*At*EPS1 in complex with the native substrate IGA, molecular docking using Autodock vina 1.1.2 (*13*) was employed to obtain an initial binding pose of IGA, which was then refined by simulated annealing using GROMACS 2020.2 (*14*). Specifically, five docking poses with the highest affinity were solvated and ionized with 0.05 M NaCl and then adopted as the starting structures for simulated annealing. Following a 1000-step energy minimization and a 1-ns NVT equilibration at 300 K, the temperature was increased to 500 K at a speed of 0.1 K/ps; after a 50-ns NPT equilibration, the temperature was reduced to 400 K at a speed of 0.05 K/ps, followed by a 5-ns NPT equilibration. The temperature was then brought to 300 K at the same speed, followed by another 5-ns NPT equilibration, and, finally, a 1000-step energy minimization. During the above simulations, protein backbones of α-helices and β-sheets were restrained with the exception of residues within 3 Å of IGA, the latter of which, along with the substrate, were allowed to move freely. The simulated annealing simulations were conducted in six replicas per docking pose. The centroid structure obtained from clustering analysis over the 30 refined structures was taken as the initial structure for the subsequent MD simulations of holo-*At*EPS1 in complex with IGA. Simulations of *At*EPS1 in complex with the yielded products (SA and NPG) and CGA were initialized by superimposing these ligands to the aforementioned refined structure of *At*EPS1 in complex with IGA.

After initialization, 2-μs MD simulations in four replicas were performed using GROMACS 2020.2 (*14*) for each system (Sup. Table. S4). These systems were placed in a dodecahedron box with a margin of at least 12 Å from any protein atoms, solvated by explicit water molecules described by the TIP3P model (*15*), and then neutralized by 0.05 M NaCl. The initial atomic velocity was assigned according to the Maxwell-Boltzmann distribution at 300 K. All systems were subjected to energy minimization, followed by a 1-ns NVT equilibration and a 1-ns NPT equilibration, with heavy atoms of the protein backbone positionally restrained. The resulting structures then were used to launch the microsecond production runs listed in Supplementary Table 4.

All simulations were performed using the CHARMM36 force field (*16*) with the ligand molecules parameterized using CHARMM General Force Field (CGenFF) (*16, 17*). Initial parameters of the ligands were obtained from the CGenFF website (*18, 19*) and then optimized using force field toolkit (FFTK) (*20*) in VMD 1.9.4 (*21*) as well as Gaussian09 (*22*). In all simulations, van der Waals interactions were smoothly switched off from 8 Å to 9 Å, while electrostatic interactions were calculated using the particle mesh Ewald (PME) method (*23*) with a cutoff of 9 Å. All systems were regulated with velocity-rescaling temperature coupling (300 K) (*24*) and Berendsen’s pressure coupling (1 bar) (*24, 25*). All bonds with H atoms were constrained using the LINCS algorithm (*26*). Clustering and other analyses were performed on all four replica trajectories of a given system. PyMOL 2.4.0 was used for visualization while atomic distances involving bound IGA were measured by VMD 1.9.4 (*21*).

---

**Supplementary Table 1.**
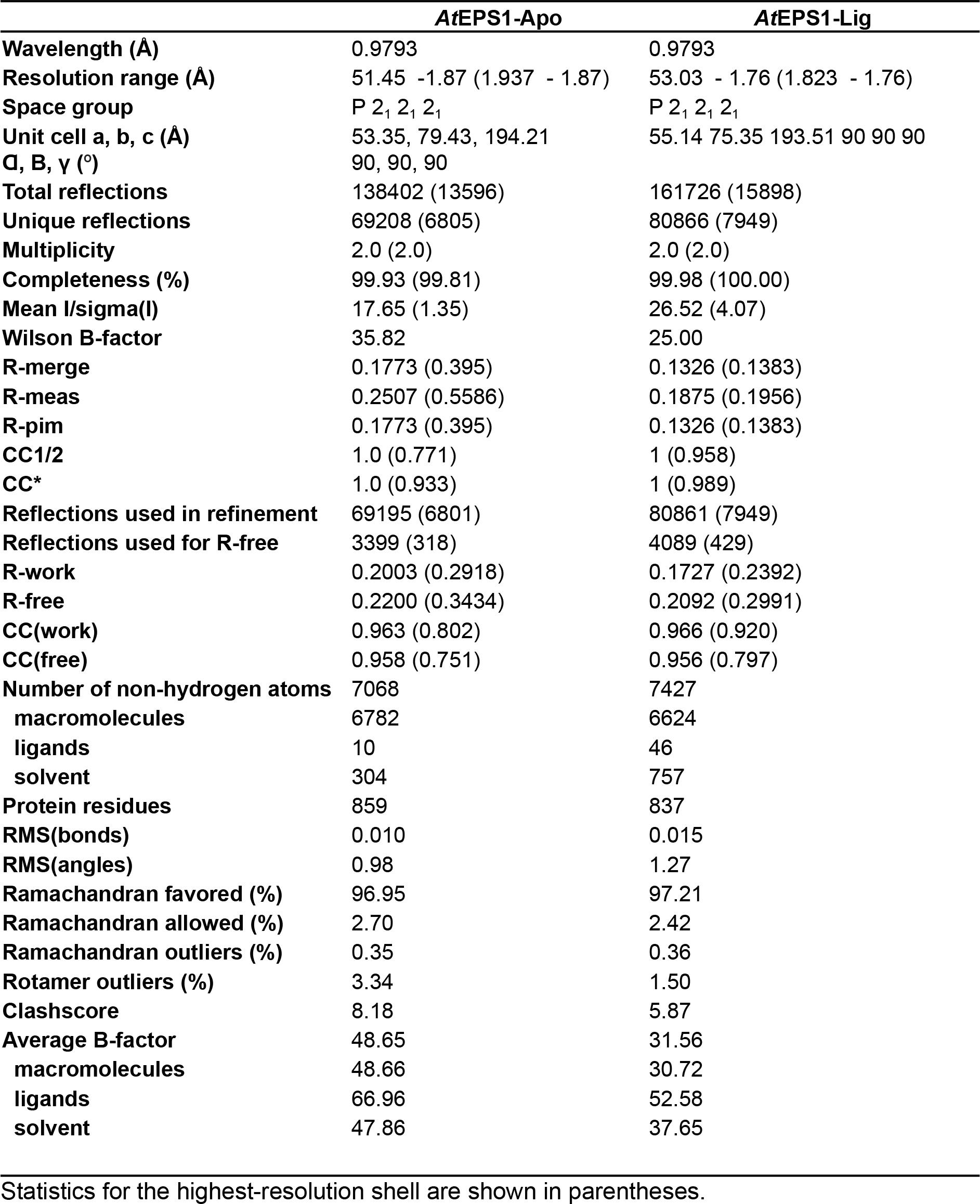
Data collection and structure refinement statistics.

**Supplementary Table 2.** BLAST E-values of various BAHDs vs *At*EPS1.

| BAHD enzymes | PDB ID | E-value |
| --- | --- | --- |
| *Selaginella moellendorffii* HCT | 6DD2 | 5.00E-37 |
| *Arabidopsis thaliana* HCT | 5KJS | 1.00E-33 |
| *Coffea canephora* HCT | 4G0B | 2.00E-31 |
| *Coleus blumei* HCT | 5KJV | 1.00E-28 |
| *Panicum virgatum* HCT | 5FAL | 2.00E-27 |
| *Sorghum bicolor* HCT | 4KE4 | 1.00E-25 |
| *Rauvolfia serpentina* VS | 2BGH | 1.00E-25 |
| *Coleus blumei* RAS | 6MK2 | 1.00E-19 |
| *Dendranthema morifolium* AT | 2E1U | 4.00E-19 |
| *Nicotiana tabacum* MAT1 | 2XR7 | 4.00E-16 |

**Supplementary Table 3.** Pairwise structural alignment Z-scores of various BAHD crystal structures vs *At*EPS1 structure.

| BAHD enzymes | PDB ID | z-score |
| --- | --- | --- |
| *Coffea canephora* HCT | 4G0B | 38.8 |
| *Coleus blumei* HCT | 5KJV | 38.6 |
| *Selaginella moellendorffii* HCT | 6DD2 | 38.6 |
| *Sorghum bicolor* HCT | 4KE4 | 37.7 |
| *Panicum virgatum* HCT | 5FAL | 37.5 |
| *Arabidopsis thaliana* HCT | 5KJS | 37.5 |
| *Nicotiana tabacum* Mat1 | 2XR7 | 36.4 |
| *Dendranthema morifolium* AT | 2E1U | 35.8 |
| *Rauvolfia serpentina* VS | 2BGH | 35.2 |
| *Coleus blumei* RAS | 6MK2 | 34.5 |

**Supplementary Table 4.** List of MD simulations conducted in this study.

| **System** | **Enzyme** | **Ligand** | **Time** |
| --- | --- | --- | --- |
| Apo-EPS1 | Apo-AtEPS1 | None | 2 μs ×<br>4 replicas |
| Holo-EPS1-CAG | Holo-AtEPS1 | (2-(3-carboxyphenoxy)acetyl)-L-glutamic acid |  |
| Holo-EPS1-IGA |  | Isochorismoyl-glutamate A |  |
| Holo-EPS1-products |  | Salicylic acid and N-pyruvoyl-L-glutamate |  |
| Holo-EPS1-CGA |  | Chorismoyl-glutamate A |  |
| IGA in water | None | Isochorismoyl-glutamate A |  |
| **Aggregated simulation time** | | | 48 μs |

**Figure S1.**
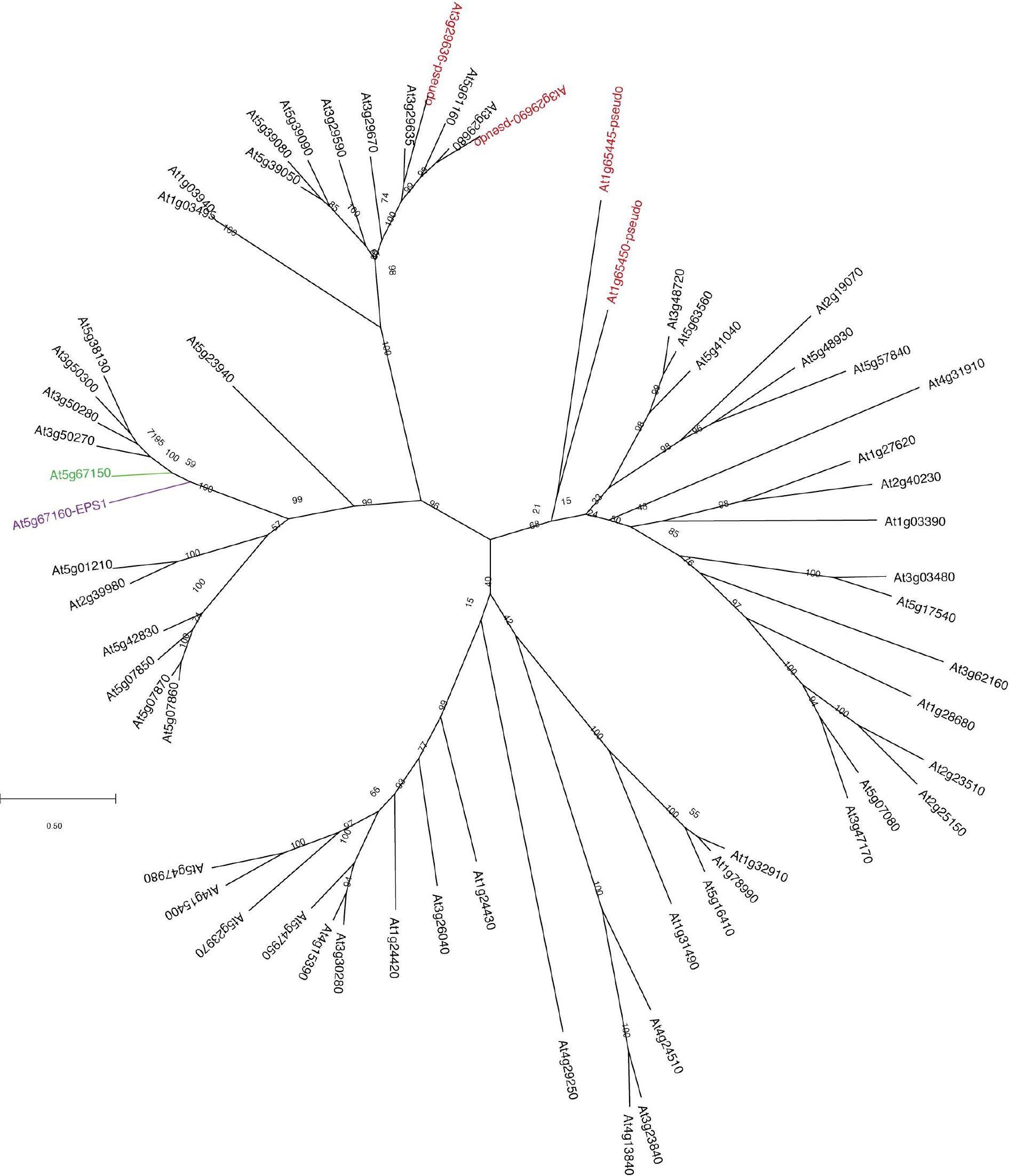
A maximum-likelihood phylogenetic tree of all previously annotated *Arabidopsis thaliana* HXXXD-type enzymes (*27*). Bootstrap values are indicated at the tree nodes. The bootstrap consensus unrooted trees were inferred from 500 replicates. The scale measures evolutionary distances in substitutions per amino acid. The purple and green text and branches correspond to *At*EPS1 (AT5G67160) and the EPS1 sister clade enzyme AT5G67170. Red text represents partial or pseudo HXXXD type sequences.

**Figure S2.**
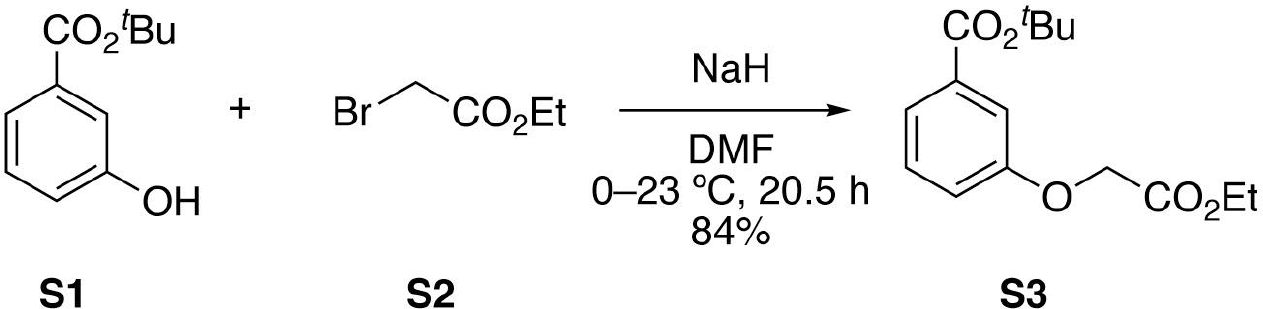
*tert*-Butyl 3-(2-Ethoxy-2-oxoethoxy)benzoate (S3). A vigorously stirred solution of phenol **S1** (500 mg, 2.58 mmol) in DMF (26.0 mL) at 0 °C was treated with NaH (60% in mineral oil) (124 mg, 3.10 mmol). The reaction mixture was warmed to 23 °C, stirred for 30 min, and then treated with bromide **S2** (0.43 mL, 3.9 mmol). The reaction mixture was stirred at 23 °C for 20 h. After 20 h, saturated aqueous NH_4_Cl (50 mL), H_2_O (100 mL), and EtOAc (150 mL) were added to the reaction mixture. The organic layer was separated and the aqueous layer was extracted with EtOAc (2 × 50 mL). The combined organic extracts were dried over Na_2_SO_4_ and concentrated on a rotary evaporator. Flash chromatography (SiO_2_, 5–10% EtOAc/hexanes) provided **S3** as a colorless film (606 mg, 84%): ^1^H NMR (CDCl_3_, 400 MHz) δ 7.63 (d, *J* = 7.7 Hz, 1H), 7.51 (s, 1H), 7.33 (t, *J* = 8.0 Hz, 1H), 7.09 (dd, *J* = 8.4, 2.7 Hz, 1H), 4.65 (s, 2H), 4.27 (q, *J* = 7.1 Hz, 2H), 1.58 (s, 9H), 1.30 (t, *J* = 7.2 Hz, 2H); ^13^C NMR (CDCl_3_, 100 MHz) δ 168.8, 165.4, 157.8, 133.6, 129.5, 123.0, 119.5, 115.0, 81.3, 65.6, 61.6, 28.3, 14.3; IR (film) ν_max_ 2978, 2932, 1758, 1711, 1586, 1486, 1440, 1368, 1294, 1193, 1163, 1105, 1083, 849, 808, 681 cm^−1^; HRMS (DART-TOF) *m/z* 280.1314 (C_15_H_20_O_5_ + H^+^ requires 281.1389).

**Figure S3.**
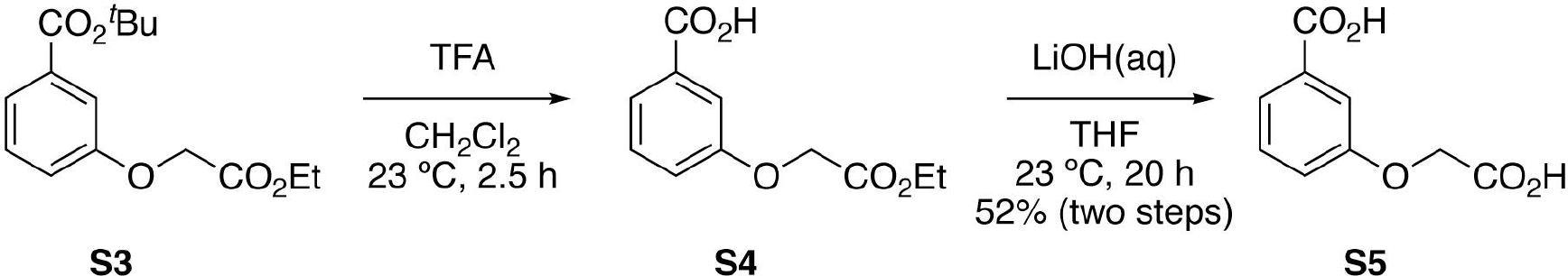
3-(Carboxymethoxy)benzoic acid (S5). A vigorously stirred solution of diester **S3** (50.0 mg, 0.178 mmol) in CH_2_Cl_2_ (900 μL) at 23 °C was treated with TFA (900 μL) and stirred for 2.5 h. After 2.5 h, the reaction mixture was concentrated and dried thoroughly under high vacuum to afford monoester **S4** which was carried forward without further purification. A solution of monoester **S4**, thus prepared, in THF (4.5 mL) at 23 °C was treated with 0.5 M aqueous LiOH (4.5 mL) and stirred vigorously for 20 h. After 20 h, the reaction mixture was concentrated under a stream of N_2_. The resulting solid residue was purified by preparative reverse-phase HPLC (CH_3_CN-0.1% TFA/H_2_O-0.1% TFA 5:95 to 95:5 over 30 min, 7 mL/min, *R*_t_ = 14.4 min) to provide diacid **S5** as a crystalline white solid (18.1 mg, 52%): mp 204–206 °C; ^1^H NMR (DMSO-*d*_6_, 400 MHz) δ 13.04 (bs, 2H), 7.55 (dt, *J* = 7.7, 1.2 Hz, 1H), 7.43–7.38 (m, 2H), 7.18 (ddd, *J* = 8.2, 2.8, 1.0 Hz, 1H), 4.74 (s, 2H); ^13^C NMR (DMSO-*d*_6_, 100 MHz) δ 170.1, 167.0, 157.8, 132.1, 129.7, 122.0, 119.4, 114.5, 64.6; IR (film) ν_max_ 3515, 3351, 3087, 1708, 1586, 1383, 1250, 1205, 1106, 1086, 834, 756, 646 cm^−1^; HRMS (DART-TOF) *m/z* 197.0443 (C_9_H_8_O_5_ + H^+^ requires 197.0450).

**Figure S4.**
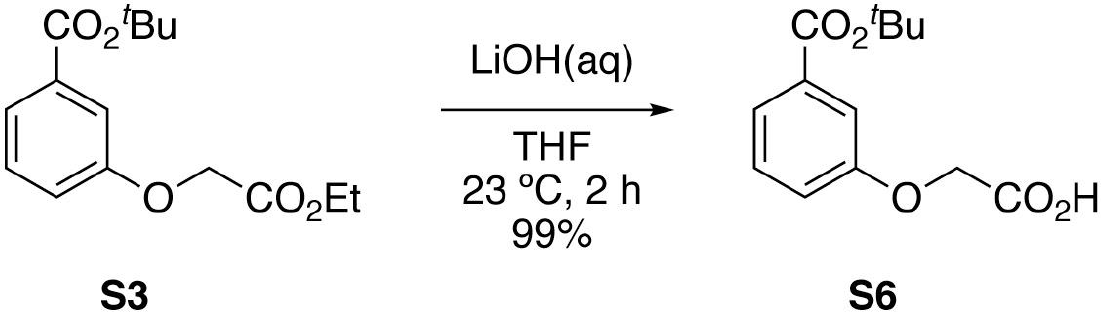
2-(3-(*tert*-Butoxycarbonyl)phenoxy)acetic acid (S6). A vigorously stirred solution of diester **S3** (250 mg, 0.892 mmol) in THF (9.0 mL) at 23 °C was treated with 0.5 M aqueous LiOH (9.0 mL). The reaction mixture was stirred at 23 °C for 2 h. After 2 h, 1 M aqueous HCl (20 mL) and CH_2_Cl_2_ (100 mL) were added to the reaction mixture. The organic layer was separated and the aqueous layer was extracted with CH_2_Cl_2_ (2 × 100 mL). The combined organic extracts were dried over Na_2_SO_4_ and concentrated on a rotary evaporator. Flash chromatography (SiO_2_, 10% MeOH-1% AcOH/CH_2_Cl_2_) provided monoester **S6** as a colorless film (223 mg, 99%): ^1^H NMR (CDCl_3_, 400 MHz) δ 10.95 (bs, 1H), 7.61 (d, *J* = 7.5 Hz, 1H), 7.52 (s, 1H), 7.28 (s, 1H), 7.08 (s, 1H), 4.64 (s, 2H), 1.58 (s, 9H); ^13^C NMR (CDCl_3_, 100 MHz) δ 173.77, 165.4, 157.4, 133.4, 129.4, 123.0, 119.4, 115.1, 81.4, 65.4, 28.1; IR (film) ν_max_ 2978, 2932, 1710, 1585, 1486, 1435, 1393, 1292, 1242, 1217, 1159, 1105, 1081, 755 cm^−1^; HRMS (DART-TOF) *m/z* 253.1106 (C_13_H_16_O_5_ + H^+^ requires 253.1076).

**Figure S5.**
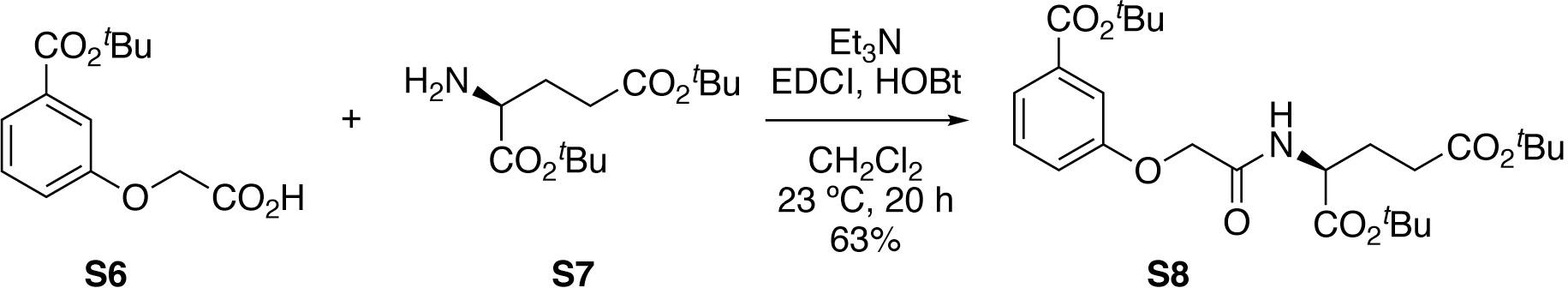
Di-*tert*-butyl (2-(3-(*tert*-Butoxycarbonyl)phenoxy)acetyl)-*L*-glutamate (S8). A vigorously stirred solution of monoester **S6** (150 mg, 0.595 mmol) in CH_2_Cl_2_ (6.0 mL) at 23 °C was treated sequentially with Et_3_N (170 μL, 1.19 mmol), protected amino acid **S7** (264 mg, 0.893 mmol), HOBt (121 mg, 0.893 mmol), and EDCI (139 mg, 0.893 mmol). The reaction mixture was stirred at 23 °C for 20 h. After 20 h, the reaction mixture was concentrated under a stream of N_2_. Flash chromatography (SiO_2_, 10–50% EtOAc/hexanes) provided amide **S8** as a colorless film (184 mg, 63%): ^1^H NMR (CDCl_3_, 400 MHz) δ 7.65 (d, *J* = 7.7 Hz, 1H), 7.54 (s, 1H), 7.36 (t, *J* = 8.0 Hz, 1H), 7.25 (bs, 1H), 7.12 (dd, *J* = 8.4, 2.7 Hz, 1H), 4.62–4.56 (m, 1H), 4.54 (s, 2H), 2.37–2.25 (m, 2H), 2.22–2.11 (m, 1H), 2.20–1.92 (m, 1H), 1.59 (s, 9H), 1.47 (s, 9H), 1.42 (s, 9H); ^13^C NMR (CDCl_3_, 100 MHz) δ 172.2, 170.7, 167.8, 165.3, 157.1, 133.9, 129.7, 123.3, 119.2, 115.4, 82.7, 81.5, 80.9, 67.4, 52.0, 31.6, 28.3, 28.2, 28.1, 27.8; IR (film) ν_max_ 2977, 2932, 1715, 1685, 1523, 1438, 1367, 1291, 1220, 1150, 846, 756 cm^−1^; HRMS (DART-TOF) *m/z* 494.2902 (C_26_H_39_NO_8_ + H^+^ requires 494.2754).

**Figure S6.**
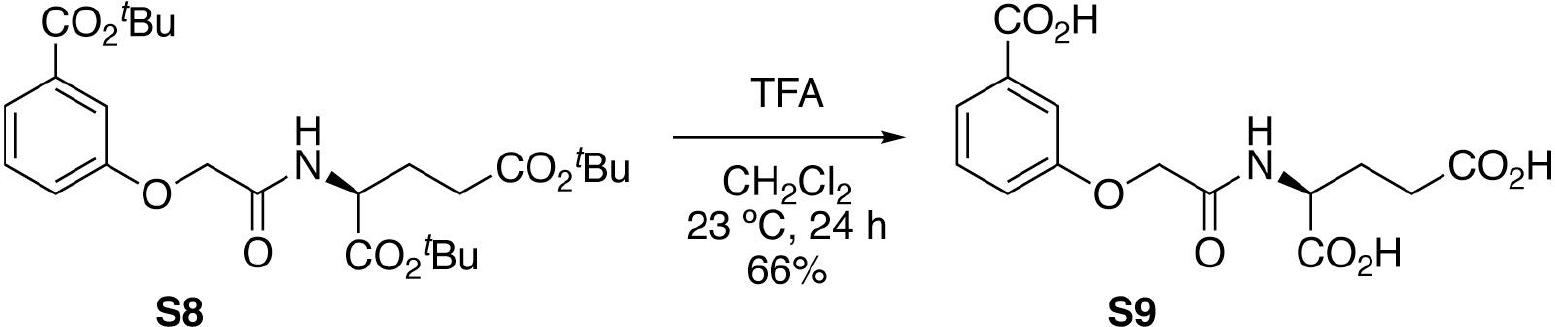
(2-(3-Carboxyphenoxy)acetyl)-*L*-glutamic acid (S9). A vigorously stirred solution of triester **S8** (100 mg, 0.203 mmol) in CH_2_Cl_2_ (1.0 mL) at 23 °C was treated with TFA (1.0 mL) and stirred for 24 h. After 2.5 h, the reaction mixture was concentrated under a stream of N_2_. The resulting residue was purified by preparative reverse-phase HPLC (CH_3_CN-0.1% TFA/H_2_O-0.1% TFA 5:95 to 95:5 over 30 min, 7 mL/min, *R*_t_ = 14.7 min) to provide triacid **S9** as an amorphous white powder (43.3 mg, 66%): mp 192–195 °C; ^1^H NMR (DMSO-*d*_6_, 400 MHz) δ 12.62 (bs, 3H), 8.38 (d, *J* = 8.0 Hz, 1H), 7.55 (d, *J* = 7.7 Hz, 1H), 7.52–7.50 (m, 1H), 7.42 (t, *J* = 7.9 Hz, 1H), 7.22 (dd, *J* = 8.2, 2.6 Hz, 1H) 4.60 (s, 2H), 4.31 (td, *J* = 9.0, 4.8 Hz, 1H), 2.26 (t, *J* = 7.5 Hz, 2H), 2.08–1.99 (m, 1H), 1.90–1.81 (m, 1H); ^13^C NMR (DMSO-*d*_6_, 100 MHz) δ 173.8, 172.9, 167.7, 167.0, 157.7, 132.2, 129.7, 122.1, 119.1, 115.5, 66.7, 51.0, 30.1, 26.1; IR (film) ν_max_ 3005, 1732, 1719, 1679, 1620, 1557, 1458, 1422, 1299, 1245, 1194, 1159, 759, 678 cm^−1^; HRMS (DART-TOF) *m/z* 326.0971 (C_14_H_15_NO_8_ + H^+^ requires 326.0876).

**Figure S7.**
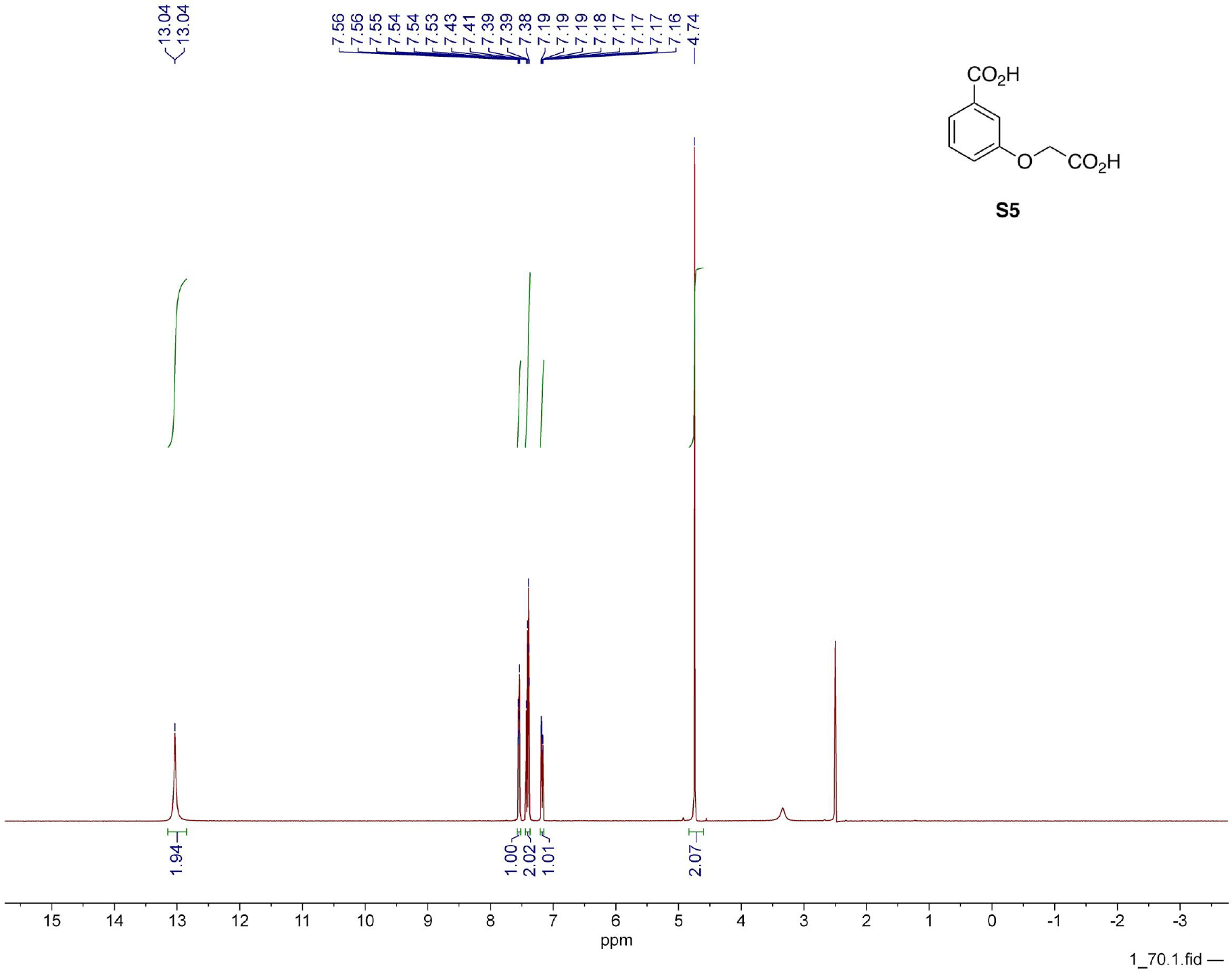
^1^H NMR spectrum of compound S5.

**Figure S8.**
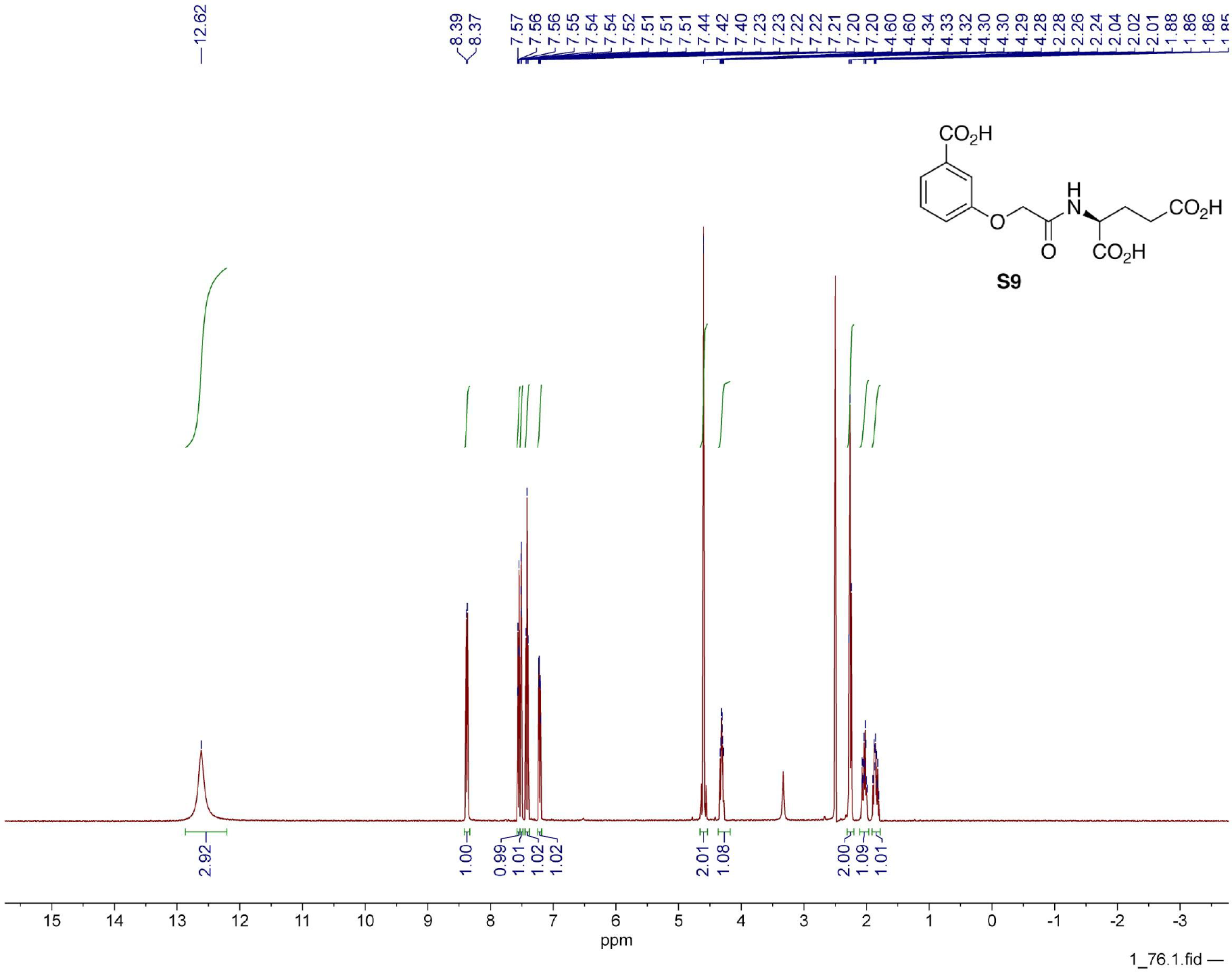
^1^H NMR spectrum of compound S9.

**Figure S9.**
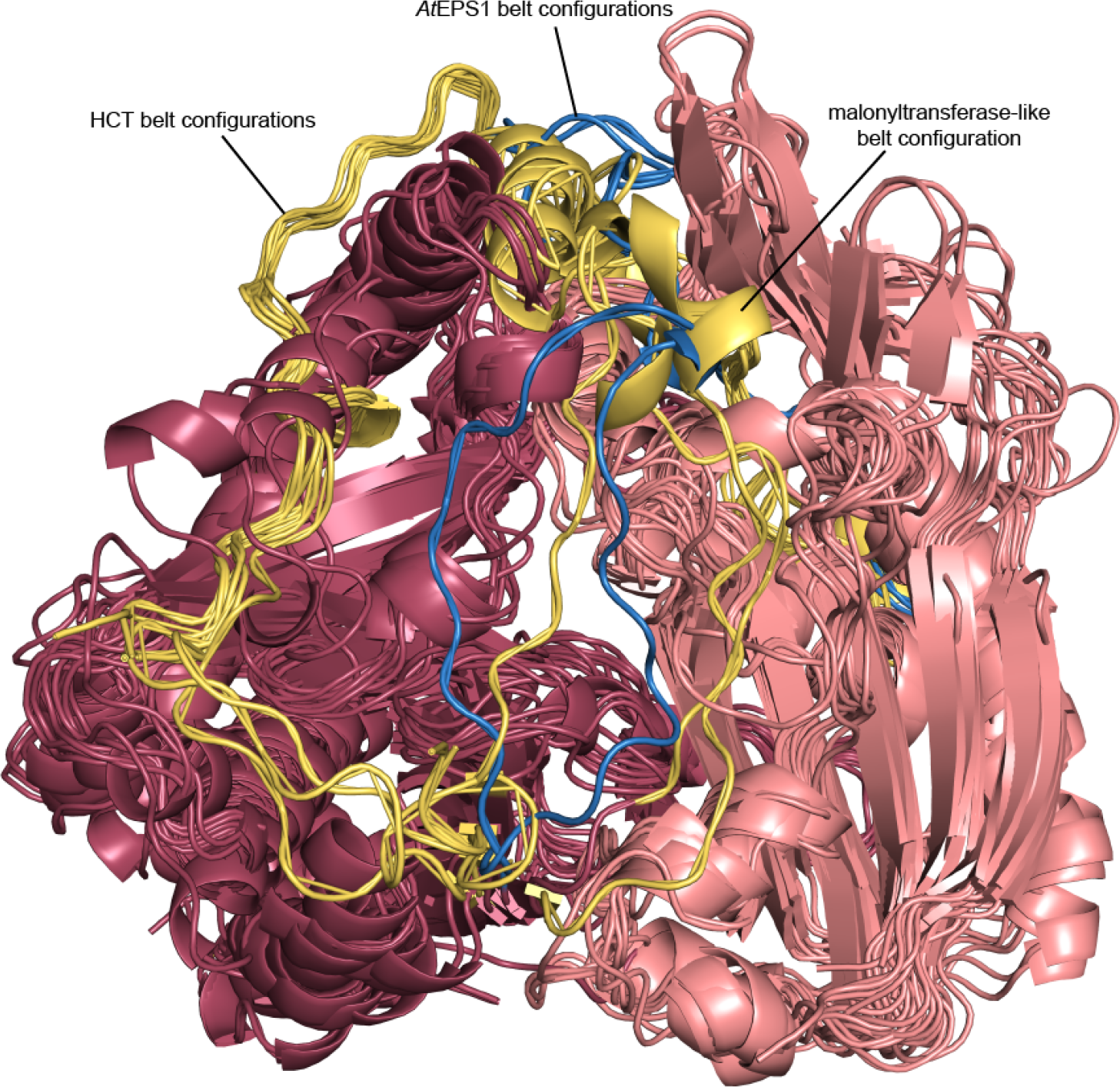
Cartoon representation of the superimposed monomers from solved plant HXXXD crystal structures (Supplementary Table 2 and 3). The N-terminal domain is displayed in salmon and the C-terminal domain is displayed in maroon. The crossover loop is displayed in slate blue in the apo and ligand bound *At*EPS structures and in yellow in the other HXXXDs. HCT structures display a movement in the belt structure relative to the orthologous Nicotiana tabacum Mat1 (2XR7), Dendranthema morifolium AT (2E1U), Rauvolfia serpentina VS (2BGH) belt structures.

**Figure S10.**
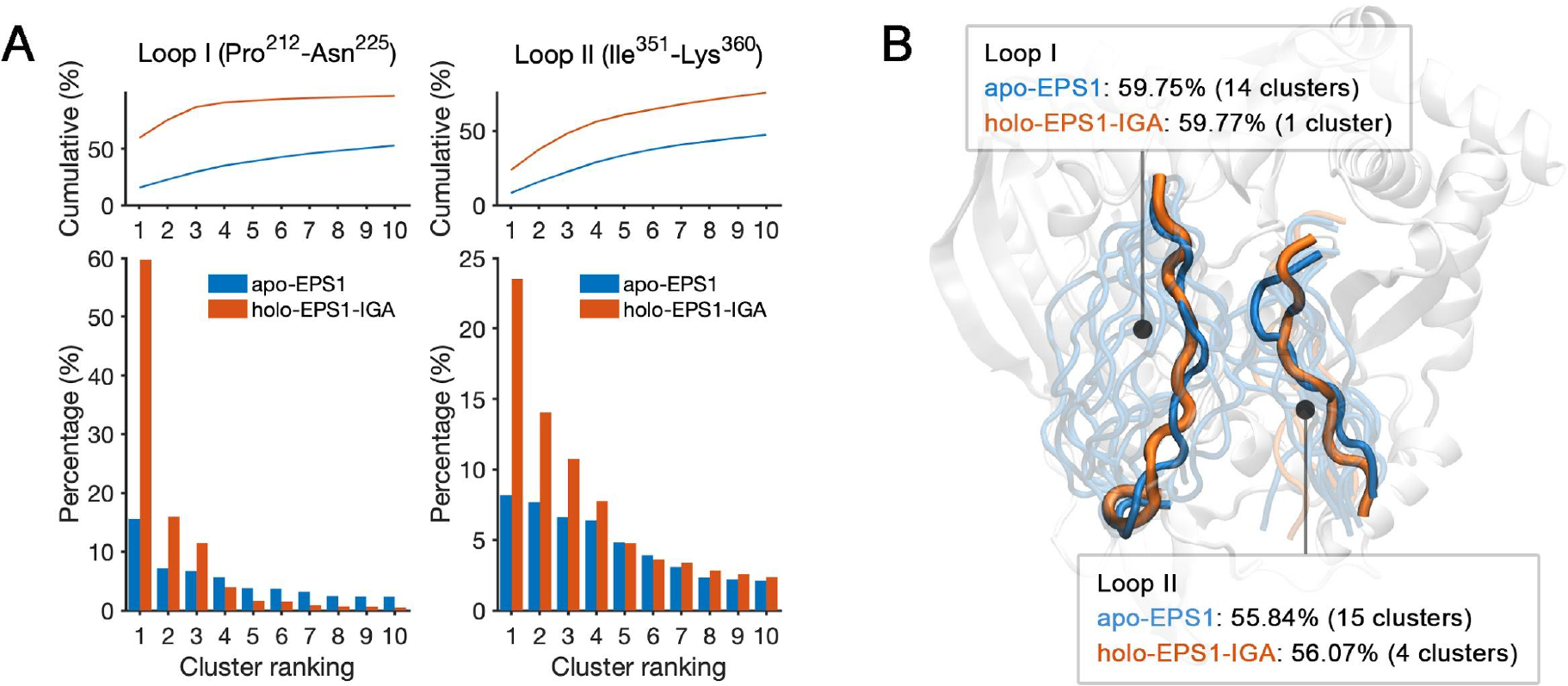
Loop dynamics in *At*EPS1. (**A**) Cumulative (top) and population (bottom) percentage of the first 10 clusters obtained from the clustering analysis of the crossover-loop (Loop I Pro^212^-Asn^225^) and lid-loop (Loop II Ile^351^-Lys^360^) in apo-EPS1 and holo-EPS1 bound with IGA over MD trajectories. (**B**) Centroid structures of the top clusters for the two loops with similar cumulative populations in the apo (blue) and holo (orange) systems. Centroid structure from the top 1 cluster of each system is displayed in an opaque manner while the rest of the centroid structures are shown in transparent representations.

**Figure S11.**
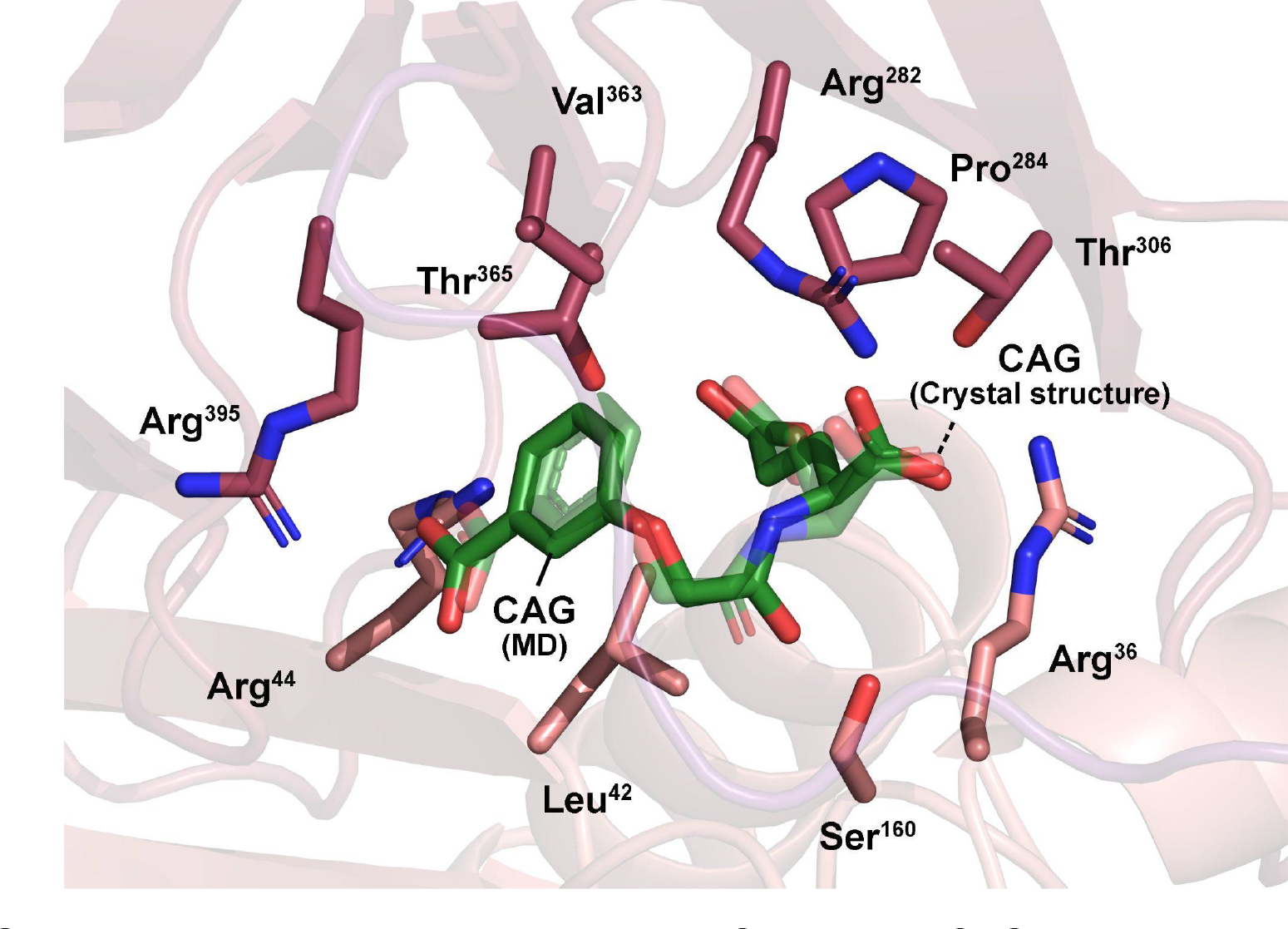
Altogether 8-μs MD simulation of *At*EPS1 bound to CAG (opaque) retains a near identical binding pose of the substrate analog to the solved crystal structure (transparent).

**Figure S12.**
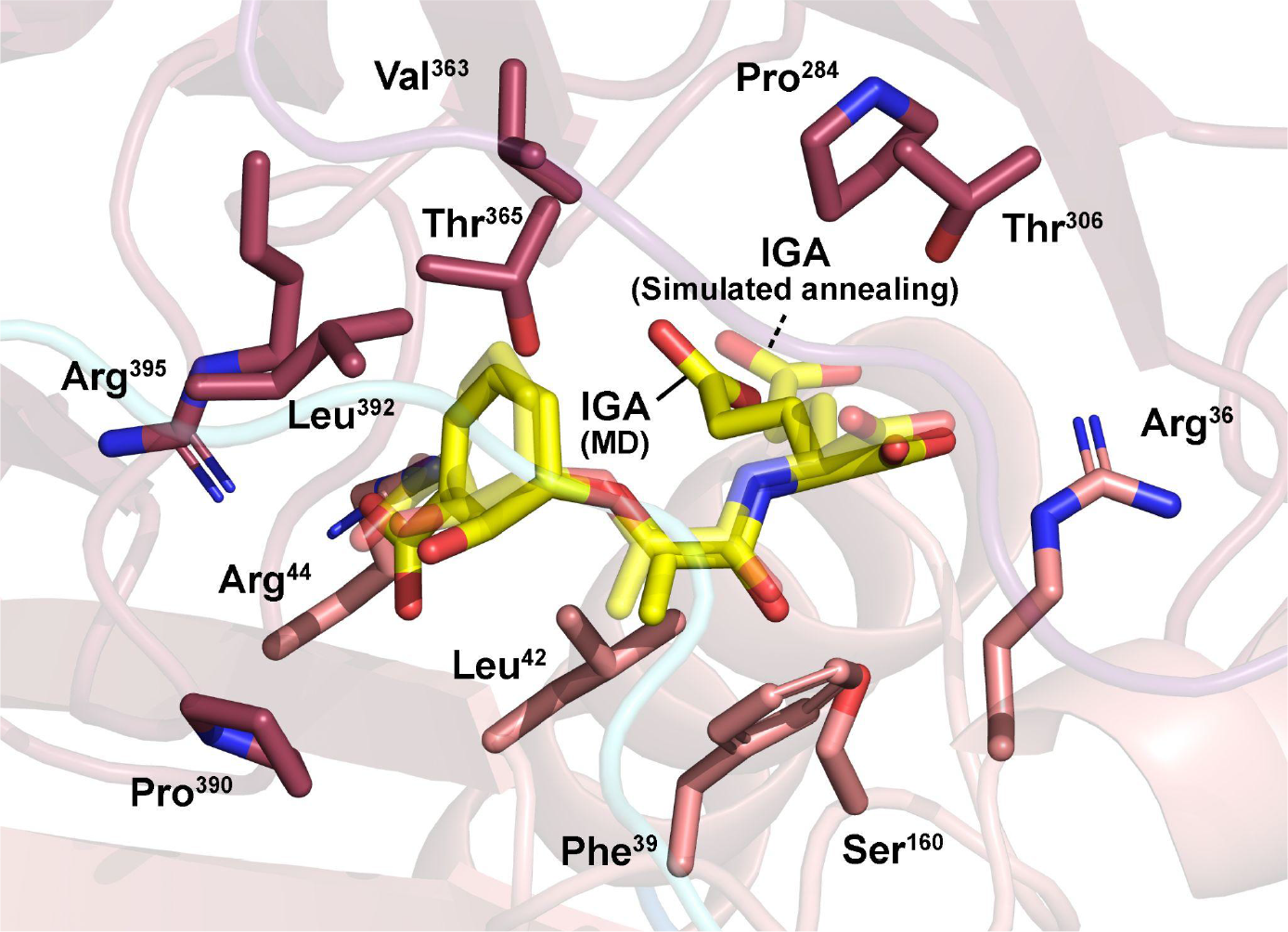
Altogether 8-μs MD simulation (opaque) and simulated annealing (transparent) reveal a similar binding pose of IGA in *At*EPS1.

**Figure S13.**
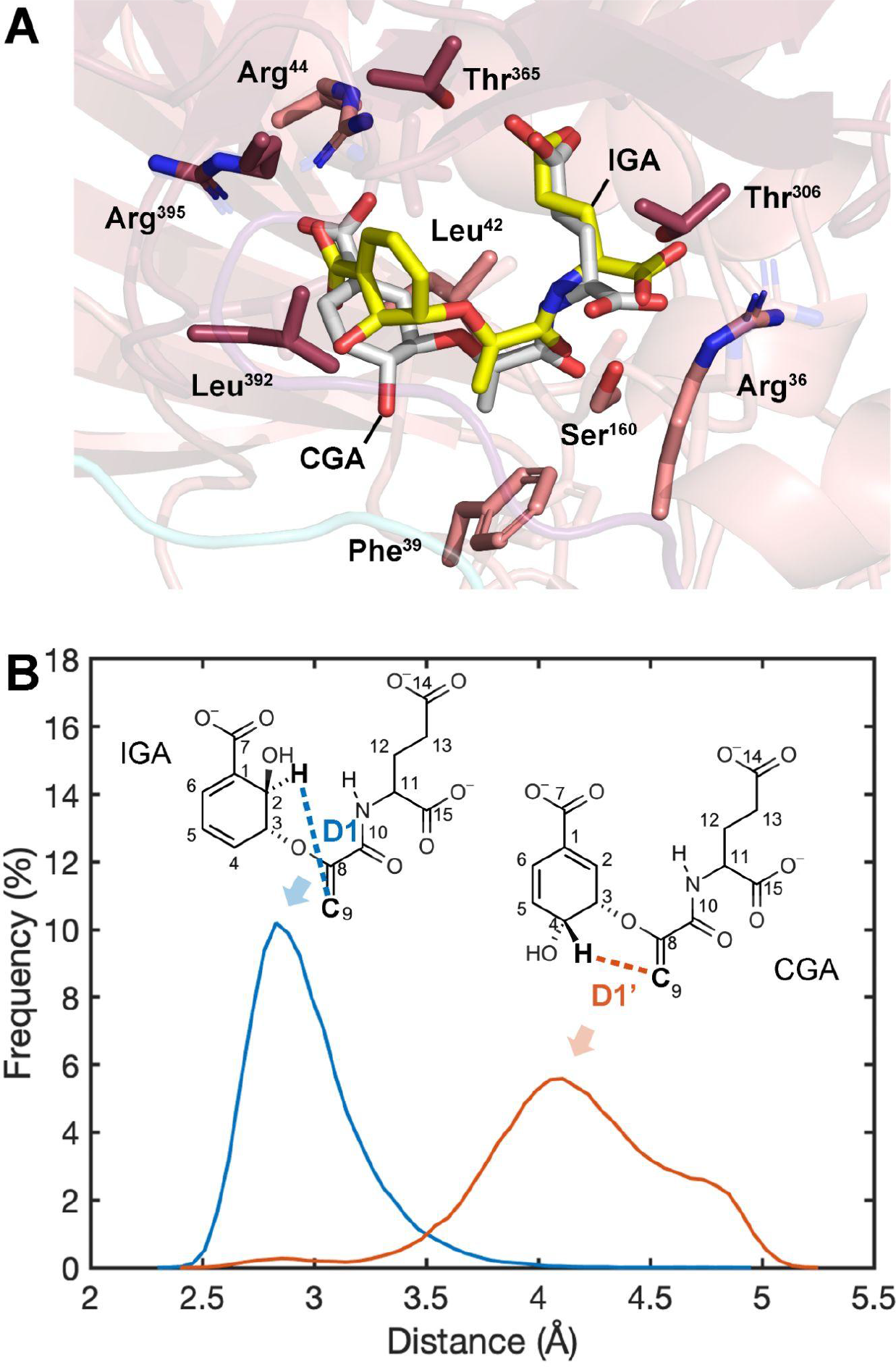
Chorismoyl-glutamate A (CGA) as a poor substrate for EPS1 (**A**) Altogether 8-μs simulations reveal that CGA (white) adopts a significantly different binding pose compared to IGA (yellow) in spite of their structural similarity. The 6-member ring of CGA adopts a pseudodiequatorial conformation and initiates hydrophobic interactions with Leu^42^. (**B**) Distance D1/D1’ measured between the transfering H and C^9^ (H^2^-C^9^ in IGA, H^4^-C^9^ in CGA). The longer D1’ distance shuts down the pericyclic reaction in CGA and stops its decomposition.

**Figure S14.**
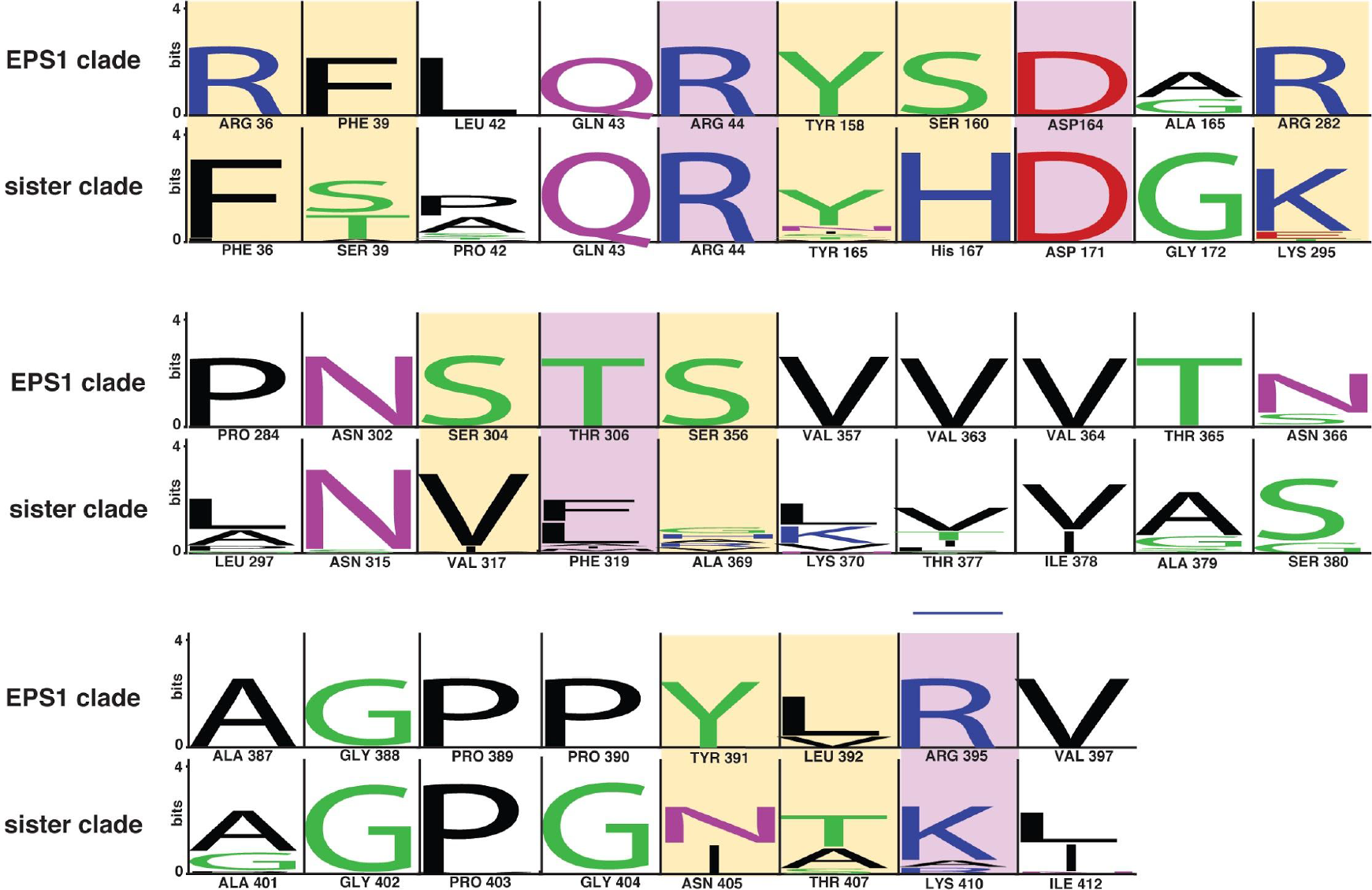
Active site pocket conservation. Select active-site-lining residues from AtEPS1 were identified and queried for conservation within the EPS1 clade (purple clade from Fig. 1B) and the sister clade (green clade from Fig.1B). The text was generated with WebLogo 3 (*3*). The height of the residue label displays the relative amino acid frequency within the selected clade. The position of the active site pocket residues from the EPS1 clade and the sister clade are referenced against the At5g67160 and At5g67160 sequences, respectively, and are listed their residue identity and position under each sequence conservation logo. Polar amino acids are colored in green, basic amino acids are colored in blue, acidic amino acids are colored in red, and hydrophobic amino acids are colored in black. Residues highlighted in purple are experimentally demonstrated to be required for IGA binding or catalysis (Fig. 4D). The yellow highlighted residues display variation between the EPS1 and sister clades of HXXXD enzymes and may play additional roles in the functional evolution of IPGL activity.

## References

1. P. Ding, Y. Ding, Stories of Salicylic Acid: A Plant Defense Hormone. Trends Plant Sci. 25, 549–565 (2020).

2. I. Raskin, Role of Salicylic Acid in Plants. Annu. Rev. Plant Physiol. Plant Mol. Biol. 43, 439–463 (1992).

3. J. León, V. Shulaev, N. Yalpani, M. A. Lawton, I. Raskin, Benzoic acid 2-hydroxylase, a soluble oxygenase from tobacco, catalyzes salicylic acid biosynthesis. Proc. Natl. Acad. Sci. U. S. A. 92, 10413–10417 (1995).

4. M. C. Wildermuth, J. Dewdney, G. Wu, F. M. Ausubel, Isochorismate synthase is required to synthesize salicylic acid for plant defence. Nature. 414, 562–565 (2001).

5. P. Silverman, M. Seskar, D. Kanter, P. Schweizer, J. P. Metraux, I. Raskin, Salicylic Acid in Rice (Biosynthesis, Conjugation, and Possible Role). Plant Physiol. 108, 633–639 (1995).

6. K. C. Chadha, S. A. Brown, Biosynthesis of phenolic acids in tomato plants infected with Agrobacterium tumefaciens. Canadian Journal of Botany. 52 (1974), pp. 2041–2047.

7. N. Yalpani, J. Leon, M. A. Lawton, I. Raskin, Pathway of Salicylic Acid Biosynthesis in Healthy and Virus-Inoculated Tobacco. Plant Physiol. 103, 315–321 (1993).

8. P. Meuwly, W. Molders, A. Buchala, J. P. Metraux, Local and Systemic Biosynthesis of Salicylic Acid in Infected Cucumber Plants. Plant Physiology. 109 (1995), pp. 1107–1114.

9. M. A. Strawn, S. K. Marr, K. Inoue, N. Inada, C. Zubieta, M. C. Wildermuth, Arabidopsis isochorismate synthase functional in pathogen-induced salicylate biosynthesis exhibits properties consistent with a role in diverse stress responses. J. Biol. Chem. 282, 5919–5933 (2007).

10. E. E. Rogers, F. M. Ausubel, Arabidopsis enhanced disease susceptibility mutants exhibit enhanced susceptibility to several bacterial pathogens and alterations in PR-1 gene expression. Plant Cell. 9, 305–316 (1997).

11. C. Nawrath, S. Heck, N. Parinthawong, J.-P. Métraux, EDS5, an Essential Component of Salicylic Acid–Dependent Signaling for Disease Resistance in Arabidopsis, Is a Member of the MATE Transporter Family. The Plant Cell. 14 (2002), pp. 275–286.

12. D. Rekhter, D. Lüdke, Y. Ding, K. Feussner, K. Zienkiewicz, V. Lipka, M. Wiermer, Y. Zhang, Feussner, Isochorismate-derived biosynthesis of the plant stress hormone salicylic acid. Science. 365, 498–502 (2019).

13. J. Mercado-Blanco, K. M. G. M. van der Drift, P. E. Olsson, J. E. Thomas-Oates, L. C. van Loon, Peter A H, Analysis of the pmsCEAB Gene Cluster Involved in Biosynthesis of Salicylic Acid and the Siderophore Pseudomonine in the Biocontrol Strain Pseudomonas fluorescensWCS374. Journal of Bacteriology. 183 (2001), pp. 1909–1920.

14. M. P. Torrens-Spence, A. Bobokalonova, V. Carballo, C. M. Glinkerman, T. Pluskal, A. Shen, J.-K. Weng, PBS3 and EPS1 Complete Salicylic Acid Biosynthesis from Isochorismate in Arabidopsis. Mol. Plant. 12, 1577–1586 (2019).

15. Z. Zheng, A. Qualley, B. Fan, N. Dudareva, Z. Chen, An important role of a BAHD acyl transferase-like protein in plant innate immunity. Plant J. 57, 1040–1053 (2009).

16. J. C. D’Auria, Acyltransferases in plants: a good time to be BAHD. Curr. Opin. Plant Biol. 9, 331–340 (2006).

17. J.-K. Weng, J. P. Noel, The remarkable pliability and promiscuity of specialized metabolism. Cold Spring Harb. Symp. Quant. Biol. 77, 309–320 (2012).

18. O. Levsh, Y.-C. Chiang, C. F. Tung, J. P. Noel, Y. Wang, J.-K. Weng, Dynamic Conformational States Dictate Selectivity toward the Native Substrate in a Substrate-Permissive Acyltransferase. Biochemistry. 55, 6314–6326 (2016).

19. L. K. Tuominen, V. E. Johnson, C.-J. Tsai, Differential phylogenetic expansions in BAHD acyltransferases across five angiosperm taxa and evidence of divergent expression among Populus paralogues. BMC Genomics. 12, 236 (2011).

20. O. Levsh, T. Pluskal, V. Carballo, A. J. Mitchell, J.-K. Weng, Independent evolution of rosmarinic acid biosynthesis in two sister families under the Lamiids clade of flowering plants. J. Biol. Chem. 294, 15193–15205 (2019).

21. Y.-C. Chiang, O. Levsh, C. K. Lam, J.-K. Weng, Y. Wang, Structural and dynamic basis of substrate permissiveness in hydroxycinnamoyltransferase (HCT). PLoS Comput. Biol. 14, e1006511 (2018).

22. A. Eudes, J. H. Pereira, S. Yogiswara, G. Wang, V. T. Benites, E. E. K. Baidoo, T. S. Lee, P. D. Adams, J. D. Keasling, D. Loqué, Exploiting the Substrate Promiscuity of Hydroxycinnamoyl-CoA:Shikimate Hydroxycinnamoyl Transferase to Reduce Lignin. Plant and Cell Physiology. 57 (2016), pp. 568–579.

23. A. M. Walker, R. P. Hayes, B. Youn, W. Vermerris, S. E. Sattler, C. Kang, Elucidation of the structure and reaction mechanism of sorghum hydroxycinnamoyltransferase and its structural relationship to other coenzyme a-dependent transferases and synthases. Plant Physiol. 162, 640–651 (2013).

24. L. A. Lallemand, C. Zubieta, S. G. Lee, Y. Wang, S. Acajjaoui, J. Timmins, S. McSweeney, J. M. Jez, J. G. McCarthy, A. A. McCarthy, A structural basis for the biosynthesis of the major chlorogenic acids found in coffee. Plant Physiol. 160, 249–260 (2012).

25. B. A. Manjasetty, X. H. Yu, S. Panjikar, G. Taguchi, M. R. Chance, C. J. Liu, Crystal Structure of Nicotiana tabacum malonyltransferase (NtMat1) complexed with malonyl-coa (2011), doi:10.2210/pdb2xr7/pdb.

26. H. Unno, F. Ichimaida, H. Suzuki, S. Takahashi, Y. Tanaka, A. Saito, T. Nishino, M. Kusunoki, T. Nakayama, Structural and mutational studies of anthocyanin malonyltransferases establish the features of BAHD enzyme catalysis. J. Biol. Chem. 282, 15812–15822 (2007).

27. X. Ma, J. Koepke, S. Panjikar, G. Fritzsch, J. Stöckigt, Crystal structure of vinorine synthase, the first representative of the BAHD superfamily. J. Biol. Chem. 280, 13576–13583 (2005).

28. C. Wang, J. Li, M. Ma, Z. Lin, W. Hu, W. Lin, P. Zhang, Structural and Biochemical Insights Into Two BAHD Acyltransferases (SHT and SDT) Involved in Phenolamide Biosynthesis. Front. Plant Sci. 11, 610118 (2020).

29. M. Ohashi, C. S. Jamieson, Y. Cai, D. Tan, D. Kanayama, M.-C. Tang, S. M. Anthony, J. V. Chari, J. S. Barber, E. Picazo, T. B. Kakule, S. Cao, N. K. Garg, J. Zhou, K. N. Houk, Y. Tang, An enzymatic Alder-ene reaction. Nature. 586, 64–69 (2020).

30. M. S. DeClue, K. K. Baldridge, D. E. Künzler, P. Kast, D. Hilvert, Isochorismate pyruvate lyase: a pericyclic reaction mechanism? J. Am. Chem. Soc. 127, 15002–15003 (2005).

31. A. L. Lamb, Pericyclic reactions catalyzed by chorismate-utilizing enzymes. Biochemistry. 50, 7476–7483 (2011).

## References

1. M. A. Larkin, G. Blackshields, N. P. Brown, R. Chenna, P. A. McGettigan, H. McWilliam, F. Valentin, I. M. Wallace, A. Wilm, R. Lopez, J. D. Thompson, T. J. Gibson, D. G. Higgins, Clustal W and Clustal X version 2.0. Bioinformatics. 23, 2947–2948 (2007).

2. S. Kumar, G. Stecher, M. Li, C. Knyaz, K. Tamura, MEGA X: Molecular Evolutionary Genetics Analysis across Computing Platforms. Mol. Biol. Evol. 35, 1547–1549 (2018).

3. G. E. Crooks, G. Hon, J.-M. Chandonia, S. E. Brenner, WebLogo: a sequence logo generator. Genome Res. 14, 1188–1190 (2004).

4. M. P. Torrens-Spence, A. Bobokalonova, V. Carballo, C. M. Glinkerman, T. Pluskal, A. Shen, J.-K. Weng, PBS3 and EPS1 Complete Salicylic Acid Biosynthesis from Isochorismate in Arabidopsis. Mol. Plant. 12, 1577–1586 (2019).

5. M. P. Torrens-Spence, T. Pluskal, F.-S. Li, V. Carballo, J.-K. Weng, Complete Pathway Elucidation and Heterologous Reconstitution of Rhodiola Salidroside Biosynthesis. Mol. Plant. 11, 205–217 (2018).

6. T. Pluskal, S. Castillo, A. Villar-Briones, M. Oresic, MZmine 2: modular framework for processing, visualizing, and analyzing mass spectrometry-based molecular profile data. BMC Bioinformatics. 11, 395 (2010).

7. J. Xia, D. S. Wishart, Using MetaboAnalyst 3.0 for Comprehensive Metabolomics Data Analysis. Curr. Protoc. Bioinformatics. 55, 14.10.1–14.10.91 (2016).

8. M. Mülbaier, A. Giannis, The Synthesis and Oxidative Properties of Polymer-Supported IBX. Angew. Chem. Int. Ed Engl. 40, 4393–4394 (2001).

9. L. A. Lallemand, C. Zubieta, S. G. Lee, Y. Wang, S. Acajjaoui, J. Timmins, S. McSweeney, J. M. Jez, J. G. McCarthy, A. A. McCarthy, A structural basis for the biosynthesis of the major chlorogenic acids found in coffee. Plant Physiol. 160, 249–260 (2012).

10. A. Vagin, A. Teplyakov, MOLREP: an Automated Program for Molecular Replacement. Journal of Applied Crystallography. 30 (1997), pp. 1022–1025.

11. G. N. Murshudov, P. Skubák, A. A. Lebedev, N. S. Pannu, R. A. Steiner, R. A. Nicholls, M. D. Winn, F. Long, A. A. Vagin, REFMAC5 for the refinement of macromolecular crystal structures. Acta Crystallogr. D Biol. Crystallogr. 67, 355–367 (2011).

12. P. Emsley, B. Lohkamp, W. G. Scott, K. Cowtan, Features and development of Coot. Acta Crystallogr. D Biol. Crystallogr. 66, 486–501 (2010).

13. O. Trott, A. J. Olson, AutoDock Vina: improving the speed and accuracy of docking with a new scoring function, efficient optimization, and multithreading. J. Comput. Chem. 31, 455–461 (2010).

14. M. J. Abraham, T. Murtola, R. Schulz, S. Páll, J. C. Smith, B. Hess, E. Lindahl, GROMACS: High performance molecular simulations through multi-level parallelism from laptops to supercomputers. SoftwareX. 1-2 (2015), pp. 19–25.

15. W. L. Jorgensen, J. Chandrasekhar, J. D. Madura, R. W. Impey, M. L. Klein, Comparison of simple potential functions for simulating liquid water. The Journal of Chemical Physics. 79 (1983), pp. 926–935.

16. R. B. Best, X. Zhu, J. Shim, P. E. M. Lopes, J. Mittal, M. Feig, A. D. Mackerell Jr, Optimization of the additive CHARMM all-atom protein force field targeting improved sampling of the backbone φ, ψ and side-chain χ(1) and χ(2) dihedral angles. J. Chem. Theory Comput. 8, 3257–3273 (2012).

17. K. Vanommeslaeghe, E. Hatcher, C. Acharya, S. Kundu, S. Zhong, J. Shim, E. Darian, O. Guvench, P. Lopes, I. Vorobyov, A. D. Mackerell, CHARMM general force field: A force field for drug-like molecules compatible with the CHARMM all-atom additive biological force fields. Journal of Computational Chemistry (2009), p. NA–NA, doi:10.1002/jcc.21367.

18. K. Vanommeslaeghe, A. D. MacKerell Jr, Automation of the CHARMM General Force Field (CGenFF) I: bond perception and atom typing. J. Chem. Inf. Model. 52, 3144–3154 (2012).

19. K. Vanommeslaeghe, E. P. Raman, A. D. MacKerell Jr, Automation of the CHARMM General Force Field (CGenFF) II: assignment of bonded parameters and partial atomic charges. J. Chem. Inf. Model. 52, 3155–3168 (2012).

20. C. G. Mayne, J. Saam, K. Schulten, E. Tajkhorshid, J. C. Gumbart, Rapid parameterization of small molecules using the Force Field Toolkit. J. Comput. Chem. 34, 2757–2770 (2013).

21. W. Humphrey, A. Dalke, K. Schulten, VMD: Visual molecular dynamics. Journal of Molecular Graphics. 14 (1996), pp. 33–38.

22. M. Frisch, F. Clemente, Gaussian 09, Revision D. 01, MJ Frisch, GW Trucks, HB Schlegel, GE Scuseria, MA Robb, JR Cheeseman, G. Scalmani, V. Barone, B. Mennucci, GA Petersson, H. Nakatsuji, M. Caricato, X. Li, HP Hratchian, AF Izmaylov, J. Bloino, G. Zhe (2009).

23. T. Darden, D. York, L. Pedersen, Particle mesh Ewald: AnN⋅log(N) method for Ewald sums in large systems. The Journal of Chemical Physics. 98 (1993), pp. 10089–10092.

24. G. Bussi, D. Donadio, M. Parrinello, Canonical sampling through velocity rescaling. J. Chem. Phys. 126, 014101 (2007).

25. R. S. Levy, C. Tendler, N. VanDevanter, P. D. Cleary, A group intervention model for individuals testing positive for HIV antibody. Am. J. Orthopsychiatry. 60, 452–459 (1990).

26. B. Hess, H. Bekker, H. J. C. Berendsen, Johannes G E, LINCS: A linear constraint solver for molecular simulations. Journal of Computational Chemistry. 18 (1997), pp. 1463–1472.

27. L. K. Tuominen, V. E. Johnson, C.-J. Tsai, Differential phylogenetic expansions in BAHD acyltransferases across five angiosperm taxa and evidence of divergent expression among Populus paralogues. BMC Genomics. 12, 236 (2011).

